# The Rho GAP RRC-1 is required for the assembly or stability of integrin adhesion complexes and is a member of the PIX pathway in muscle

**DOI:** 10.1101/2022.07.02.498541

**Authors:** Jasmine C. Moody, Hiroshi Qadota, Guy M. Benian

## Abstract

GTPases cycle between active GTP bound and inactive GDP bound forms. Exchange of GDP for GTP is catalyzed by guanine nucleotide exchange factors (GEFs). GTPase activating proteins (GAPs) accelerate GTP hydrolysis, to promote the GDP bound form. Recently, we reported that the GEF called PIX is required for assembly or stability of integrin adhesion complexes (IAC) in striated muscle. A GAP for the PIX pathway had not been identified in any cell type or organism. A screen in *C. elegans* of mutants in 18 proteins containing Rho GAP domains and expressed in muscle revealed that loss of function of *rrc-1* results in loss of IAC components at the muscle cell boundary (MCB). RRC-1 contains an SH3 domain and a Rho GAP domain, and is localized to the IACs of MCBs, like PIX-1. *rrc-1* mutants show reduced accumulation of IAC components at the MCB, sarcomere disorganization, and reduced whole animal locomotion. Knockdown of *git-1*, which encodes a PIX scaffold protein, reduces the level of RRC-1. Localization at MCBs of RRC-1 depends on *pix-1*, and the localization of PIX-1 depends on *rrc-1*. These results suggest that RRC-1 is a RhoGAP for the PIX pathway in muscle.

## Introduction

Integrin adhesion complexes (IAC), also known as focal adhesions, consist of the transmembrane heterodimeric proteins α/β integrin as well as hundreds of other proteins that are associated both from the extracellular matrix (ECM) and especially intracellularly (Anthis and Campbell, 2011; Sun et al., 2014; Bachir et al., 2014; Horton et al., 2015). IACs are important for many cell types. The adhesion of cells to a matrix is crucial for both tissue formation and for cell migration. In stationary cells like muscle, these complexes are rather stable, but in motile cells they are dynamic, with new complexes assembled at the leading edge and older complexes disassembled at the trailing edge (Anthis and Campbell, 2011). When integrins are expressed on the cell surface they are in a compact or bent or inactive state, unable to bind to their ECM targets, but can become activated to bind via several triggers (e.g. chemokines, local increase in P(4,5)IP_2_) that lead to binding of the cytoplasmic tail of β-integrin to talin. Binding to talin results in integrin assuming a more open conformation, able to bind to extracellular targets (Tadokoro et al., 2003). Kindlin is also involved in integrin activation by clustering of talin-activated integrins, at least in platelets (Ye et al., 2013). Although we understand the steps involved in the formation of IACs, we do not know how the composition of an IAC is determined or regulated, and we do not know what determines where an IAC forms, how many IACs form, and what their spacing will be.

In striated muscle, which includes both skeletal and cardiac muscle, myofibrils at the periphery of the cell are attached to the cell membrane and ECM via “costameres”(Ervasti, 2003; Henderson et al., 2017), muscle-specific IACs. Costameres are involved in anchorage of the muscle cell to the ECM, and transmission of force. *C. elegans* is an outstanding genetic model organism in which to learn new principles about muscle (Gieseler et al., 2017). The major striated muscle is found in the body wall and is used for locomotion (Benian and Epstein, 2011). Similar to striated muscle in other animals, the thin filaments are attached to Z-disk like structure (dense bodies), and the thick filaments are attached to M-lines. The sarcomeres are restricted to a narrow ∼1.5 mm zone adjacent to the cell membrane along the outer side of the muscle cell, and all the dense bodies and M-lines are anchored to the muscle cell membrane and ECM. The base of dense bodies and M-lines contain integrin adhesion complexes (IACs) and much is known about their protein composition (Gieseler et al., 2017). Additional IACs are located at the muscle cell boundaries, where they form attachment plaques that anchor the muscle cell to a thin layer of ECM that lies between adjacent muscle cells (Qadota et al., 2017). Thus, in *C. elegans* muscle IACs are located at M-lines, dense bodies and muscle cell boundaries (MCBs), and although the base of each consists of integrins and a set of core proteins, they also contain proteins specific for each site (Gieseler et al., 2017). IACs at MCBs consist of a subset of proteins that are found at dense bodies (Qadota et al., 2017).

Recently, we reported that a protein in *C. elegans*, PIX-1 (orthologous to β-PIX in mammals), is required for the assembly or stability of IACs at MCBs (Moody et al., 2020). A PIX signaling pathway is important for the mammalian nervous (Schmalxigaug et al., 2009; Ramakers et al., 2012; Huang et al., 2011) and immune systems (Volinsky et al., 2006; Missy et al., 2008), and for the control of distal tip cell shape and migration (important for formation of the germline) (Lucanic and Cheng, 2008), and for epithelial morphogenesis (Zhang et al., 2011) in *C. elegans*. However, no prior study had demonstrated a function for PIX in striated muscle in any organism. PIX proteins contain an SH3 domain, and a Rho GEF domain that activates the small GTPases Rac and Cdc42. In *C. elegans*, PIX-1 is localized to the IACs present at the muscle boundaries and the IACs at M-lines and dense bodies. As compared to wild type, a *pix-1* null mutant shows reduced levels of activated, GTP-bound Rac in muscle. Interestingly, either deficiency or overexpression of PIX-1 results in disrupted MCBs and decreased nematode muscle function, suggesting that the level of PIX-1 needs to be tightly controlled. The Rho GEF domain of PIX proteins promote the exchange of GDP for GTP, thus converting inactive to active Rac or Cdc42. In the PIX pathway, the active GTP bound Rac or Cdc42 binds to and activates a PAK family protein kinase, which then phosphorylates one or more unknown substrates to somehow promote assembly of IACs.

Rho family GTPases (Rho, Rac, Cdc42) cycle between active (GTP-bound), and inactive (GDP-bound) states. Activation occurs via guanine-nucleotide exchange factors (GEFs)(e.g. PIX) that promote exchange of GDP with GTP, and inactivation occurs via GTPase-activating proteins (GAPs), which promote the hydrolysis of GTP to GDP. Perhaps because of the cycling requirement, the terminal phenotypes of loss of function for a GEF and loss of function for a GAP (for a particular GTPase and cellular function), are often the same. For example, in yeast, loss of function of the GTPase Bud1p (similar to mammalian Rap GTPases) has a similar phenotype to loss of function of its GEF, Bud5p, and its GAP, Bud2p (Michelitch and Chant, 1996). In *C. elegans*, the same embryonic cytokinesis defect is found for loss of function for *rho- 1* (RhoA), *rga-3* (Rho GAP), *rga-4* (Rho GAP), and *ect-2* (Rho GEF)(Morita et al., 2005; Schonegg et al., 2007; Jantsch-Plunger et al., 2000; Canman et al., 2008). For the PIX pathway in *C. elegans*, the GEF is PIX-1, and the GAP is unknown. In fact, a GAP for the PIX pathway has not been reported for any organism. We hypothesized that for the PIX pathway in nematode muscle, the loss of function for the GEF, PIX-1, and an unknown GAP, would be the same. Using an easily scorable phenotype (i.e. loss of IAC components at the MCB), we screened mutants in genes predicted to encode Rho GAP proteins, and identified one protein, RRC-1, which is required for assembly or stability of IACs at MCBs. RRC-1 contains an SH3 domain and a Rho GAP domain. We found that RRC-1 is localized to the IACs of MCBs, like PIX-1. Loss of function mutants of *rrc-1* show reduced accumulation of multiple IAC components at the MCB and reduced whole animal locomotion, but in addition show sarcomere disorganization. Knockdown of *git-1*, which encodes the PIX scaffold protein GIT, reduces the level of RRC-1. Moreover, the localization at MCBs of RRC-1 depends on *pix-1*, and the localization of PIX-1 depends on *rrc-1*. These results suggest that RRC-1 is a RhoGAP for the PIX-1 pathway in muscle.

## Results

### Either decreased or increased activity of the PIX-1 pathway results in disorganization of IACs at MCBs

The output of the PIX signaling pathway in mammals and nematodes is that p21-activated kinases (PAKs) which are serine/threonine protein kinases are activated by binding to GTP-bound Rac or Cdc42. Because activation of PAK requires that a GEF first activates Rac or Cdc42, and inactivation occurs by a GAP, we hypothesized that a PAK-1 kinase dead and a PAK-1 kinase constitutively active mutant might have the same phenotype. We have an easily scorable muscle phenotype for the status of the PIX-1 pathway, i.e. deficiency of any component results in disorganization of integrin adhesion complexes (IAC) at the muscle cell boundary (MCB), including a deficiency of *pak-1* (Moody et al., 2020). We used CRISPR/Cas9 to generate mutant worms carrying either a catalytically dead or constitutively activating mutations for *pak-1*. For the catalytically-dead mutant, we replaced K324 with A. In nearly all protein kinases, a K at this position is found in the small lobe of the kinase domain and coordinates ATP and helps transfer γ-phosphate. Mutation of this K to A or several other amino acids inactivates most known kinases (Iyer et al., 2005).

In the absence of GTP-Rac or GTP-Cdc42, PAKs exist in a closed conformation due to binding of an N-terminal (67-150) autoinhibitory domain (AID) with the more C-terminal kinase catalytic domain (Zenke et al. 1999). Binding of GTP-Rac or Cdc42 to this AID leads to an opening of the PAK structure and activation of its phosphotransferase activity. A constitutively-active human PAK has been generated by substituting the highly conserved L107 to F in this AID (Brown et al., 1996). In the nematode protein, the homologous residue is L99, and this was mutated to F as described under Methods. As shown in Figure 1, the catalytically dead mutant, *pak-1(syb632)* which has a K324A mutation, and a constitutively active mutant, *pak-1(syb647)* which has a L99F mutation, each have a similar defect at the MCBs. Specifically, each mutant shows less accumulation of PAT-6 and increased spacing between adjacent muscle cells. This result demonstrates that either decreased or increased activity of the PIX-1 pathway can have the same terminal phenotype. Thus, we hypothesized that increased activity of the PIX-1 pathway by inactivation of a Rho GAP would have the same phenotype as decreased activity of the PIX-1 pathway by inactivation of its Rho GEF, PIX-1.

**Figure 1.**
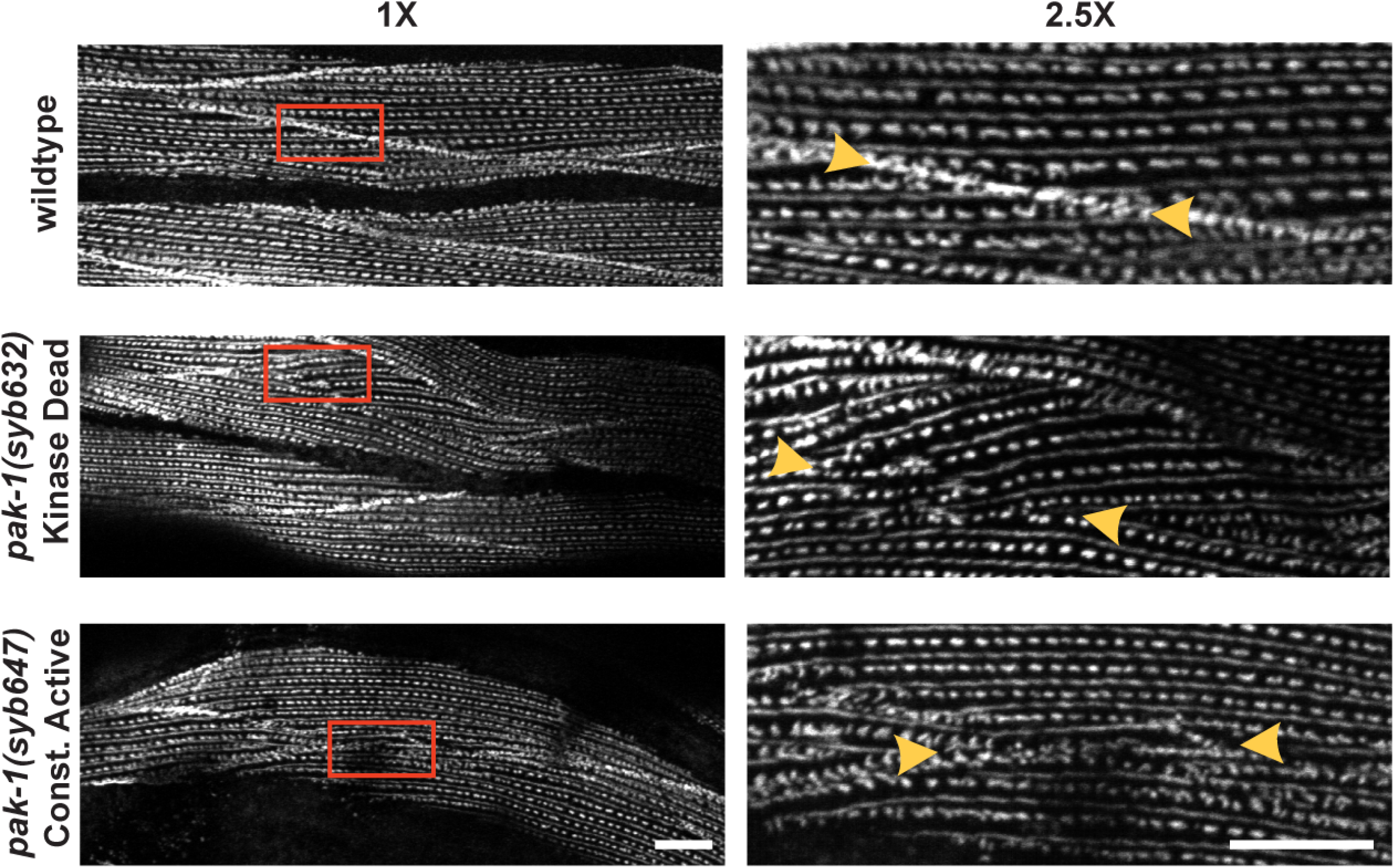
Either catalytically dead PAK-1 or constitutively active PAK-1 kinase results in a muscle cell boundary defect. Confocal microcopy imaging of wildtype, *pak-1(syb632)[K324A]* kinase dead, and *pak-1(syb647) [L99F]* constitutively active kinase mutants immunostained with anti-PAT-6 (α-parvin), shown at 1X and 2.5X optical zoom. 2.5X zoom areas are the insets indicated by the red boxes in 1X zoom images. Yellow arrows indicate muscle cell boundaries. In wild type, these structures are well-formed and “tight”, but in the *pak-1* mutants they are disrupted or missing. Scale bar, 10 μm.

### Screening for a Rho GAP for the PIX pathway

Our homology search has revealed that there are 32 genes in *C. elegans* that encode proteins harboring Rho GAP domains. Of these 32, 18 of them are expressed in body wall muscle based on SAGE data (Meissner et al., 2009). We obtained mutants for all 18 genes from the Caenorhabditis Genetics Center (see Methods) and screened them for the MCB defect using anti-PAT-6 (α-parvin) immunostaining. Mutants for two genes, *hum-7* and *rrc-1* (Supplementary Figure 1, Supplementary Table 1) each demonstrated defects of the MCB. *hum-7* encodes a 1880 residue protein containing an RA domain, a class IX myosin motor domain, 2 IQ domains, and a Rho GAP domain, and will be described elsewhere. The *rrc-1* gene encodes an approximately 750 residue protein that contains a Rho GAP domain and an SH3 (Figure 2a). Because RRC-1 is a simpler protein, we decided to focus our efforts on RRC-1. Alternative splicing of *rrc-1* produces three protein isoforms containing both domains and of approximately the same size (742-759 aa)(Figure 2b). A Rho GTPase effector pull-down assay of nematode RRC-1 expressed in mammalian tissue culture cells demonstrates that RRC-1 has GAP activity towards mammalian Rac and Cdc42 but not RhoA (Delawary et al. 2007).

**Figure 2.**
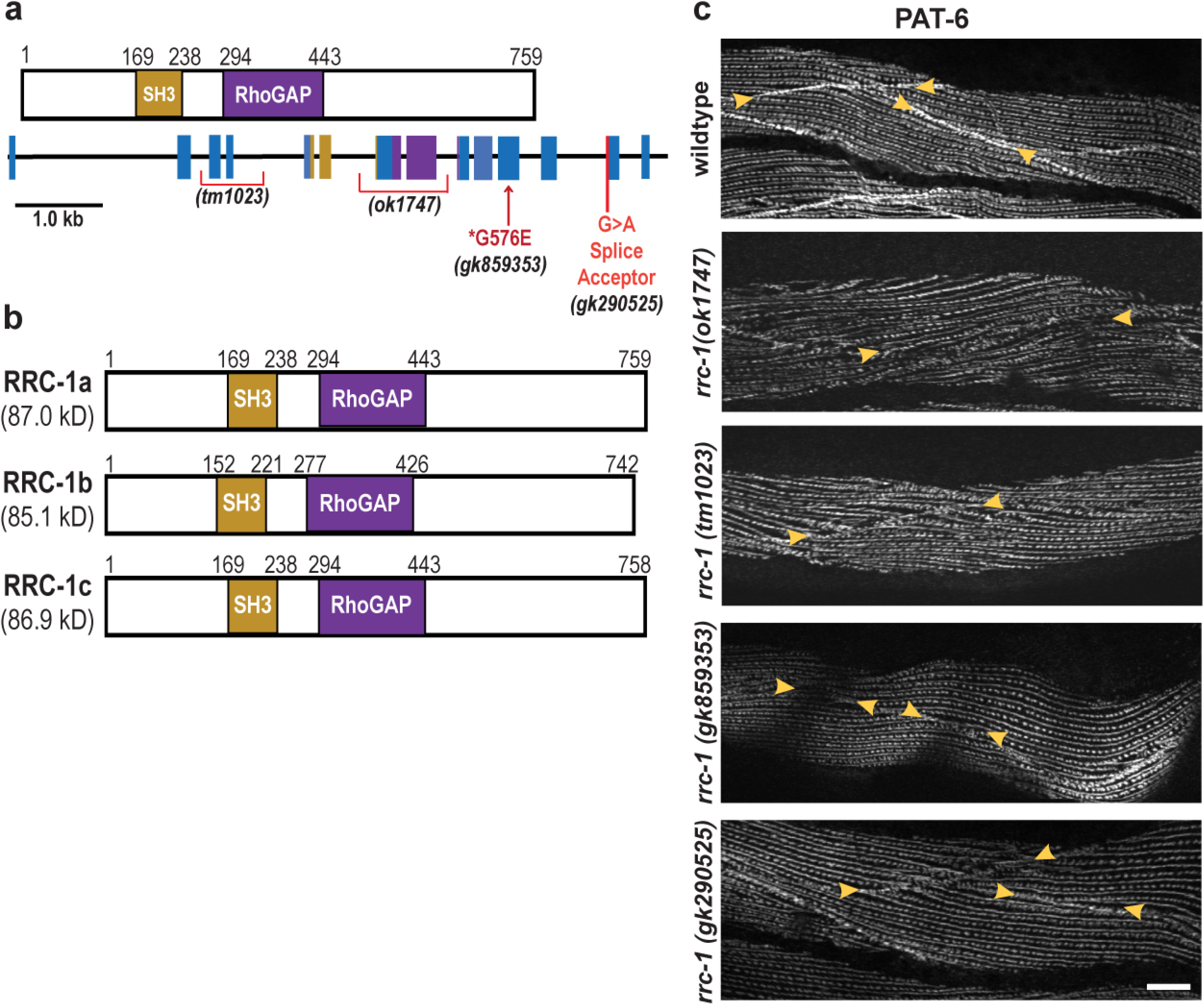
Loss of function *rrc-1* mutants show a lack of or disorganization of PAT-6 at muscle cell boundaries. **a.)** Schematic representation of domains in *C. elegans* RRC-1 isoform A, and the location and nature of the four *rrc-1* mutants within a map of the exon-intron organization of the *rrc-1* gene. **b.)** Schematic showing domain organization of the three predicted RRC-1 isoforms, generated by alternative splicing. **c.)** Confocal microscopy imaging of body wall muscle cells immunostained with antibodies to PAT-6 (α-parvin) from wildtype, two RRC-1 out-of-frame deletion allele mutants (*rrc-1(ok1747)* and *rrc-1(tm1023)),* one missense mutation, *rrc-1(gk859353)*, and one splice site mutation, *rrc-1(gk290525),* each outcrossed 5x to wildtype. Arrowheads point to the boundaries between muscle cells. Note that there is also disorganization of PAT-6 localization at M-lines and dense bodies in the deletion alleles, *ok1747* and *tm1023*. Scale bars, 10µm.

### RRC-1 is orthologous to human ARHGAP32 and ARHGAP33 proteins

Wormbase considers RRC-1 to be orthologous to three human proteins, ARHGAP31, 32 and 33, and the website notes that the best BLASTP match is to ARHGAP32. We obtained these human sequences and performed a PFAM prediction of domains and aligned and compared each of the domains to those in RRC-1a. As shown in Figure 3, all four proteins contain a RhoGAP domain. Human ARHGAP31 only contains a RhoGAP domain, whereas RRC-1a and human ARHGAP32 and 33 also contain SH3 domains. Furthermore, human ARHGAP32 and 33 have PX domains, and ARHGAP32 has an additional RPEL repeat domain. Based on the number of shared domains and their percent identities, we consider the closest human orthologs to RRC-1 to be ARHGAP32 and ARHGAP33, and probably ARHGAP33 is closer. Inspection of The Human Protein Atlas shows that all three human proteins are expressed in multiple organs, but ARHGAP31 is not expressed in skeletal or heart muscle, whereas ARHGAP32 and 33 show “medium levels” of expression in skeletal and heart muscle. These expression patterns are consistent with ARHGAP32 and 33 being closer human orthologs for RRC-1.

**Figure 3.**
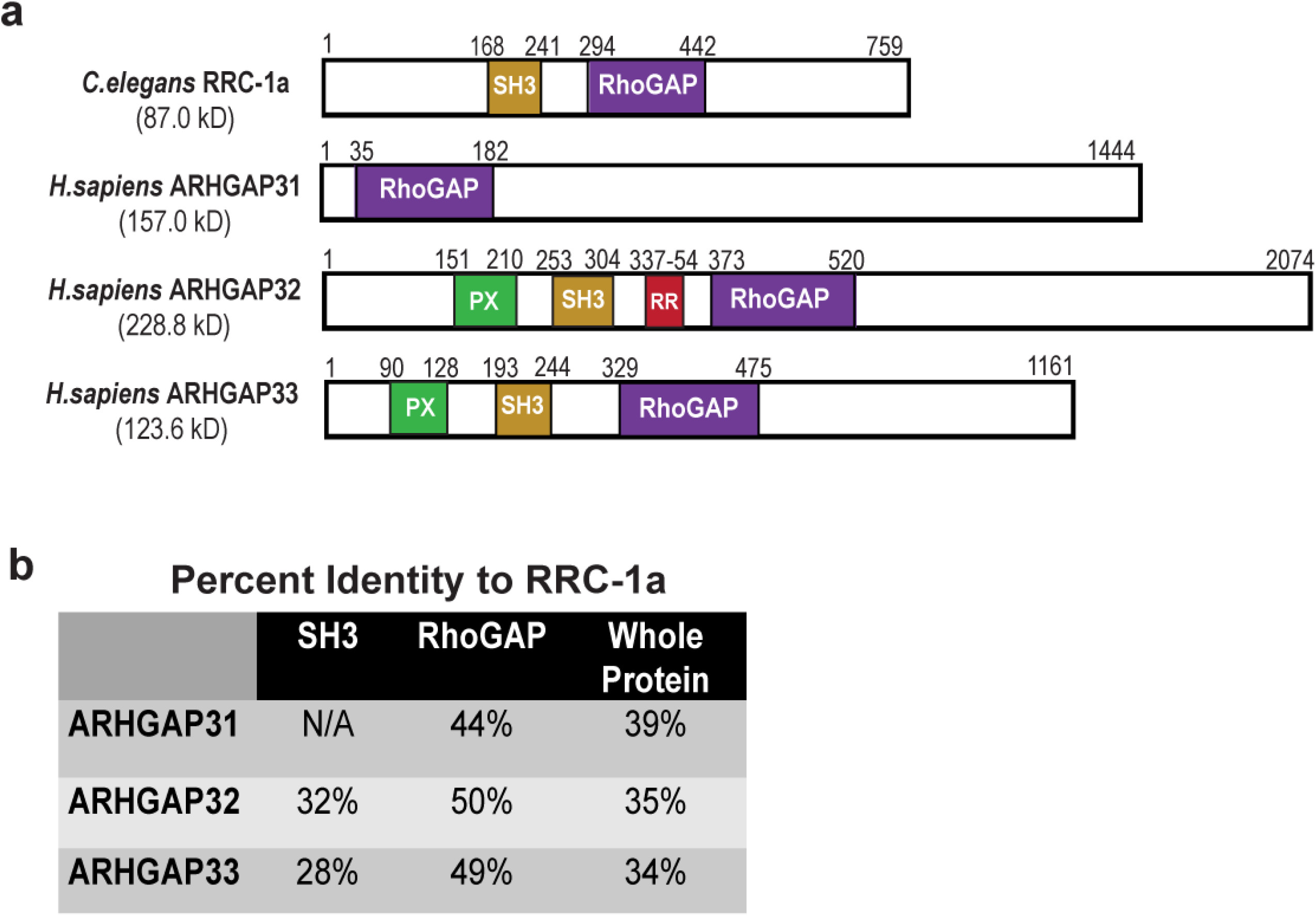
RRC-1 is orthologous to human ARHGAP32 and ARHGAP33 proteins. **a.)** Schematic representation of domain organization in C. elegans RRC-1a, and human ARHGAP31, 32 and 33 proteins, as predicted PFAM. Numbers above the long rectangles represent amino acid positions. **b.)** Table showing percent identity of each whole protein and predicted domains of the human proteins compared to nematode RRC-1a. Sequence alignment was made using NCBI pBLAST.

### *rrc-1* mutants display mis-localized or missing IAC components at muscle cell boundaries, and disorganized M-lines and dense bodies

The original allele that we characterized, *rrc-1(ok1747)*, is a frame-shifting deletion that removes exons 7 and 8 (Figure 2a). We obtained 3 more *rrc-1* mutant alleles from the Caenorhabditis Genetics Center, the first being *tm1023*, which is also a frame-shifting deletion removing exons 3 and 4. Inspection of the Million Mutation Project collection (Thompson et al., 2013) revealed 16 *rrc-1* mutants, one being a splicing defect, and 15 being missense mutations. We ordered the splicing defect mutant, and the four missense mutants that have nonconservative amino acid changes. Unfortunately, two of these strains were too difficult to grow, and one had background mutations in *pak-1* and in *unc-89,* which would confound our analysis, and thus were not pursued. Therefore, we had a collection of four *rrc-1* mutant alleles, including two deletions, one missense mutant, and one splicing acceptor mutant (Figure 2a). We outcrossed each mutant to wild type five times, to remove most of the background mutations. Figure 2c shows results from immunostaining of the four *rrc-1* alleles with antibodies to PAT-6 (α-parvin) to visualize IACs at the MCBs. All four alleles show defects, but the deletion alleles, *ok1747* and *tm1023*, are more severely affected, showing not only weak concentration of PAT-6 at the boundaries but also what appear to be gaps between adjacent cells.

By immunostaining, we found that in the deletion mutant, *rrc-1(ok1747)*, other IAC components are missing or mis-localized at MCBs (Figure 4). These IAC components include UNC-52 (perlecan) in the ECM, UNC-95 and UNC-112 (kindlin). We conclude that the Rho GAP RRC-1 is required for the assembly or stability of IACs at the MCB.

**Figure 4.**
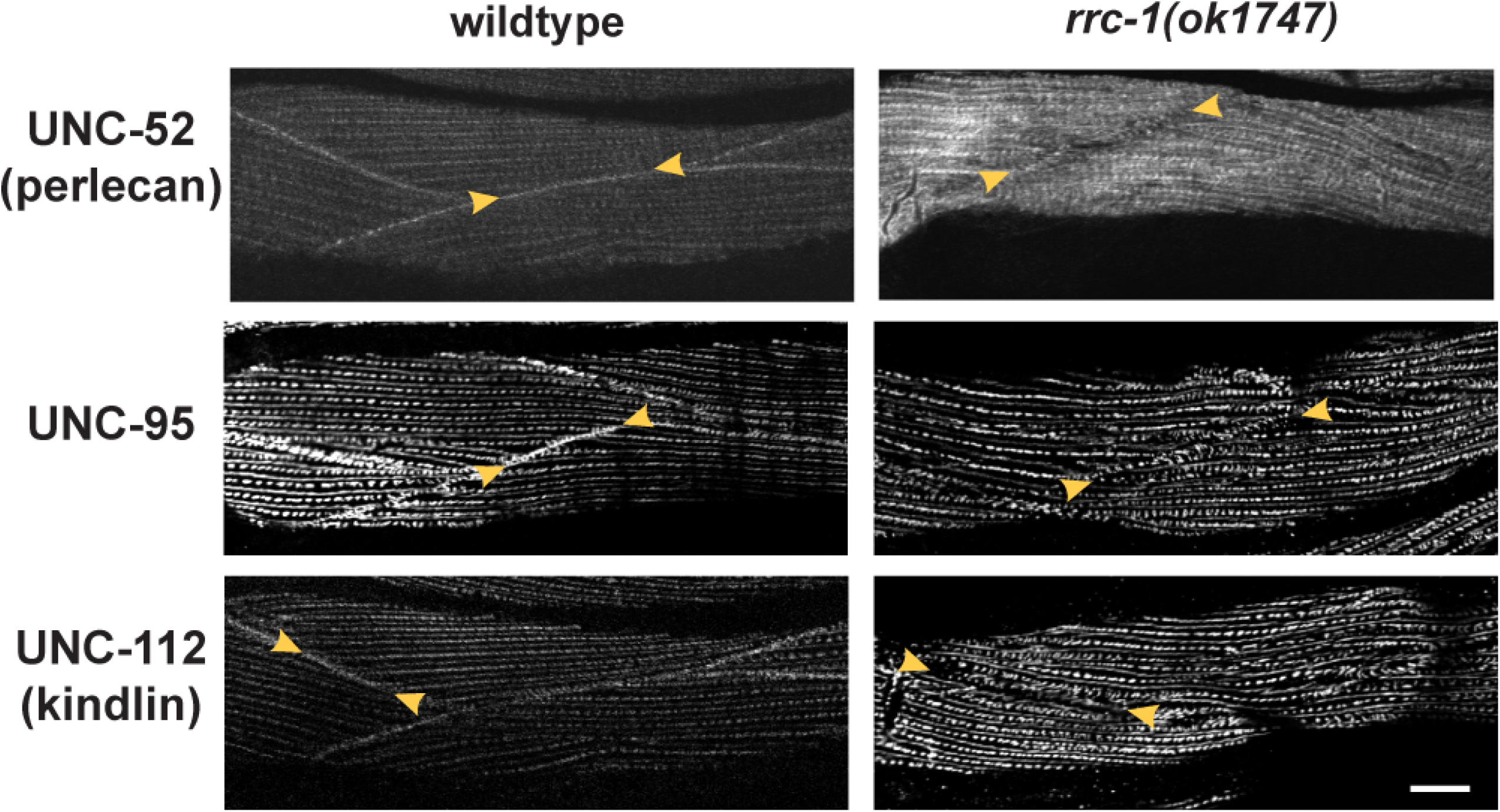
Mis-localization of multiple IAC components at the muscle cell boundaries, M-lines and dense bodies in *rrc-1* mutants. Comparison of wild type vs. *rrc-1(ok1747),* outcrossed 5x to wildtype, immunostained with antibodies to the indicated integrin adhesion complex (IAC) proteins and imaged by confocal microscopy. In parentheses are the names of the mammalian orthologs. Arrowheads denote muscle cell boundaries. Note that all three proteins are present at the muscle cell boundaries in wildtype but are missing or less tightly organized at the muscle cell boundaries in the *rrc-1* deletion mutant. In addition, there is mis-localization of each of these IAC components at M-lines and dense bodies. Each image is a representative image obtained from at least 2 fixation and immunostaining experiments, and imaging of at least three different animals. Scale bars, 10µm.

In addition, the M-lines and dense bodies are disorganized in *rrc-1(ok1747)*, especially seen upon immunostaining of UNC-95 and UNC-112 (Figure 4). The dense bodies are not as regularly punctate, sometimes being elongated, and the distance between adjacent rows of dense bodies and M-lines is increased and irregular. Thus, RRC-1 is also required for the assembly or stability of IACs at M-lines and dense bodies.

To rule out that the defects that we have observed in *rrc-1* mutants by immunostaining are artefacts resulting from incomplete fixation, we localized UNC-112-GFP in live animals using a transgenic strain. As shown in Supplementary Figure 2, UNC-112-GFP is localized to the MCBs in wild type but missing at the MCBs in *rrc-1(ok1747)*. Moreover, UNC-112-GFP is also missing at MCBs in *pix-1(gk299374)*, which allowed us to first describe the MCB boundary defect phenotype by immunostaining (Moody et al., 2020).

### *rrc-1* mutants also display disorganization of sarcomeres

Because in *rrc-1(ok1747)*, we found disorganization in M-lines, which normally cross-link thick filaments, and dense bodies, which normally attach to thin filaments, we examined the organization of the sarcomere further. We used phalloidin staining to determine the organization of thin filaments, and antibodies to myosin MHC A for the organization of thick filaments, to UNC-89 for the organization of M-lines (full depth), and to ATN-1 (α-actinin) for the organization of the main portion of dense bodies. As shown in Figure 5, all sarcomere structures are abnormal in *rrc-1(ok1747)* and *rrc-1(tm1023)*, as compared to wild type. The greatest degree of disorganization is observed in UNC-89 (M-lines), and MHC-A, which is the myosin isoform located in the middle of thick filaments where they are cross linked at the M-line. Less disorganization is observed in thin filaments (phalloidin), or with the main portions of dense bodies (ATN-1). Because similar sarcomere defects were observed in two independently generated *rrc-1* mutants that were extensively outcrossed, these defects can be attributed to loss of function of rrc-1 rather than any closely linked background mutations.

**Figure 5.**
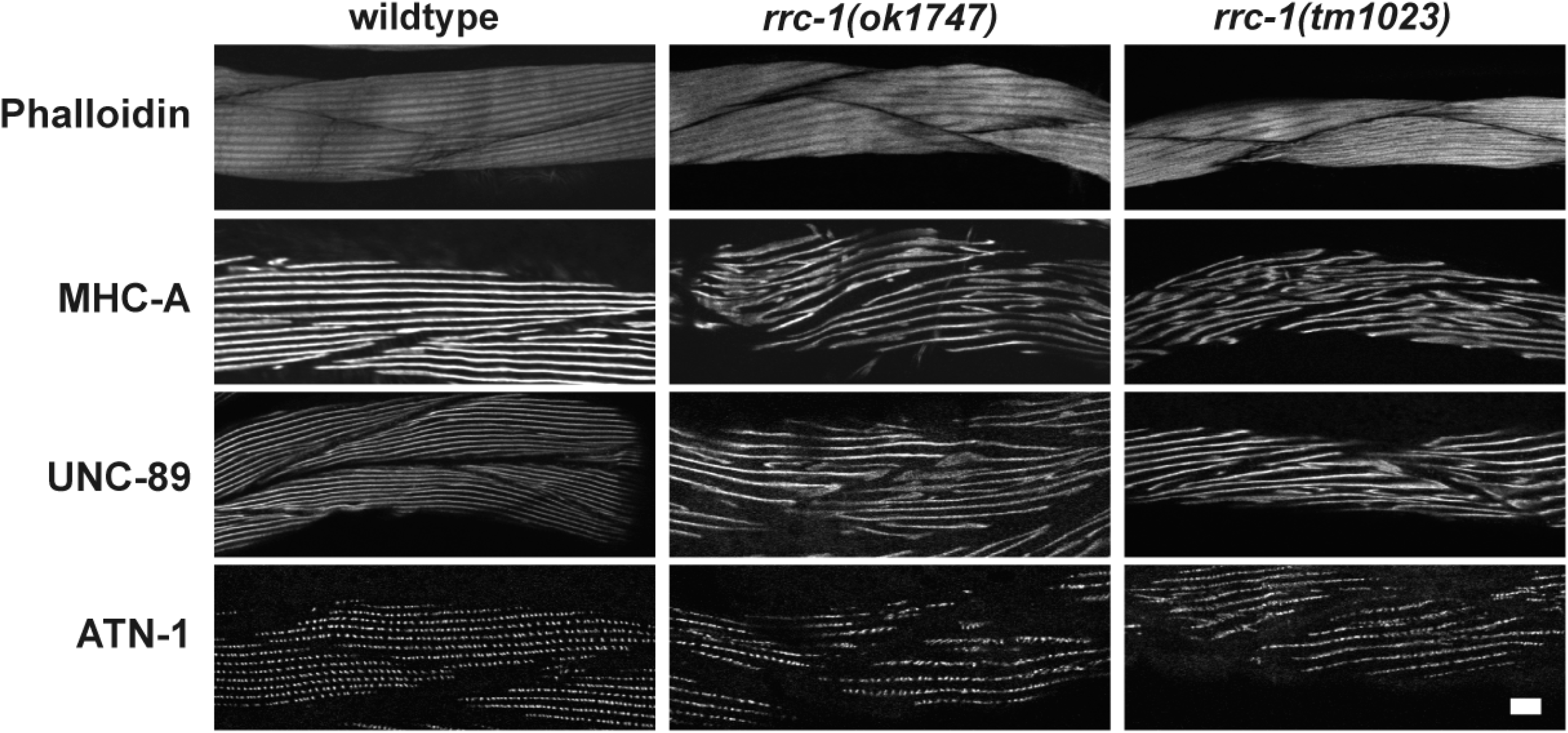
*rrc-1* mutants have disorganized sarcomeres. Confocal images of wild type, *rrc- 1(ok1747)* and *rrc-1(tm1023)* reacted with phalloidin-rhodamine (thin filaments), and antibodies to sarcomere proteins MHC A (thick filaments), UNC-89 (M-lines) and ATN-1 (dense bodies). Note the disorganization of each sarcomere component compared to wild type. Most severely affected are the thick filaments or A-bands, and the M-lines. Each image is a representative image obtained from at least two immunostaining experiments. Scale bar, 10 μm.

### *rrc-1* mutants are defective in whole animal locomotion

In *C. elegans*, the force of body wall muscle contraction that bends the worm and thus permits locomotion of the animal, is generated in the sarcomeres and transmitted through all three IAC sites, including the M-lines, the dense bodies and the adhesion plaques at the MCBs. We previously reported that *pix-1* mutants have reduced whole animal locomotion, and this is likely attributable to them having poorly organized adhesion plaques at MCBs, because they have normally organized sarcomeres, M-lines and dense bodies (Moody et al., 2020). However, *rrc-1* mutants show more extensive defects—in addition to the MCB, the M-lines, dense bodies and sarcomeres are disorganized. Thus, we conducted worm motility assays. As shown in Figure 6a, both deletion alleles, *ok1747* and *tm1023*, and the splicing mutant, *gk290525*, display reduced swimming when compared to wild type. However, the missense mutant, *gk859353*, displays swimming that is not significantly different from wild type. Although *gk859353* is a non-conservative G to E change, it resides outside of a recognizable domain and thus may not have a critical function. Crawling may be a more stringent test of the ability of a worm to move because it is likely that the worm needs to overcome the surface tension lying between itself and the agar surface. Thus, as shown in Figure 6b, all four *rrc-1* mutants exhibit reduced crawling as compared to wild type. Moreover, for all four alleles, the trends in both swimming and crawling are similar, with the deletion alleles and splicing mutant showing slower movement than the missense mutant.

**Figure 6.**
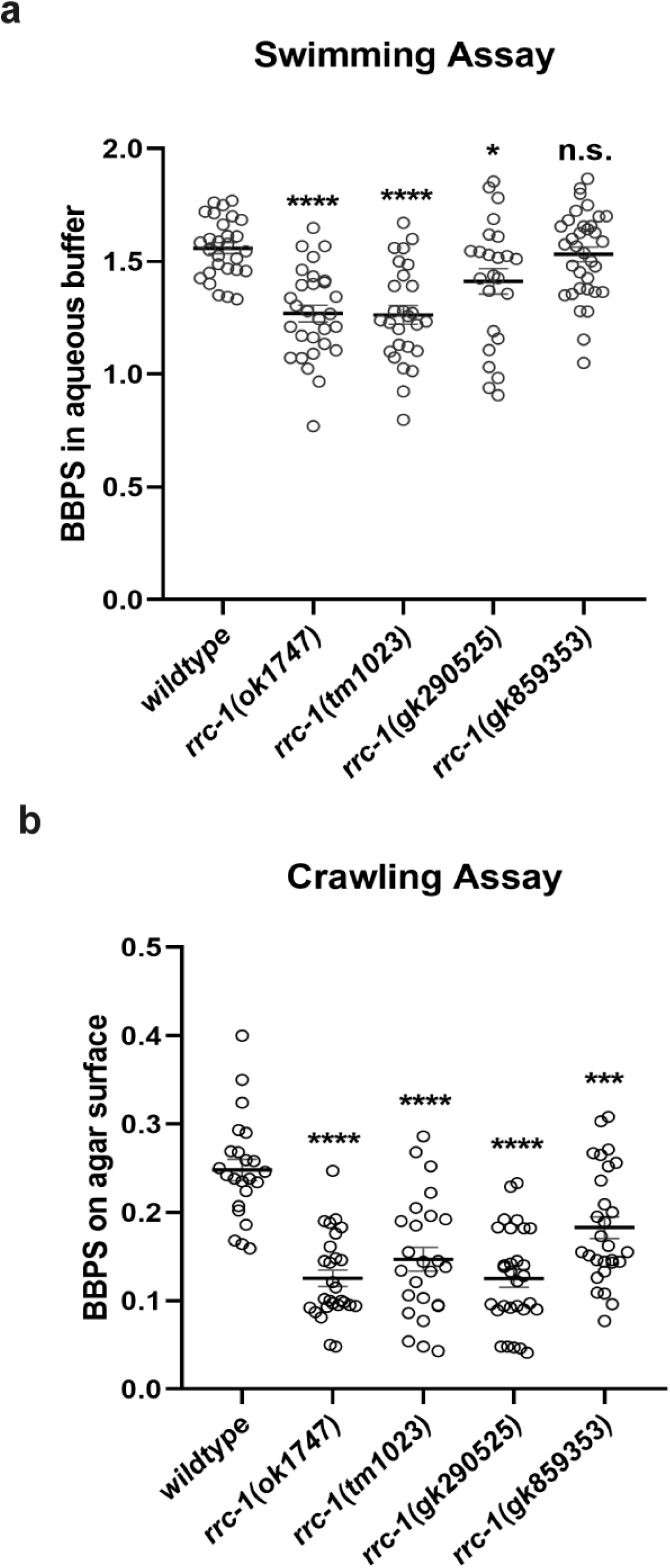
Loss of function *rrc-1* mutants have reduced whole animal locomotion. **a.)** Swimming and **b.)** crawling assays show that RRC-1 deletion mutants *rrc-1(ok1747)* and *rrc- 1(tm1023),* a splice acceptor site mutant, *rrc-1(gk290525)*, and a missense mutant, *rrc- 1(gk859353),* outcrossed 5x to wildtype result in reduced locomotion as compared to wildtype animals. Body bends per second (BBPS) are quantified for individual animals of each strain. In the graphs, each open circle represents the result from an independently selected animal. The exact n values vary, but n ≥ 23. Welch’s t-test was used to test for significance. Error bars indicate SEM, *p≤ .05, ****p≤ .0001.

### The localization of RRC-1 protein in muscle

We first attempted to make antibodies to RRC-1. Unfortunately, using two different immunogens, we failed to generate specific antibodies in rabbits. Therefore, to localize RRC-1 in muscle, we used CRISPR/Cas9 to create a worm strain, *rrc-1(syb4499)*, in which the endogenous *rrc-1* gene expresses RRC-1 with an HA tag fused to its C-terminus. As shown in Figure 7a, by western blot using anti-HA antibodies, we detect an RRC-1-HA fusion of expected size (approximately 90 kDa) from this strain but not from wild type. To test whether the HA tag might interfere with the normal function of RRC-1, we conducted locomotion assays and immunostaining of sarcomeres. As shown in Supplementary Figure 3, we observed no difference in swimming or crawling motility assays between wild type and the RRC-1::HA strain. In addition, as shown in Supplementary Figure 4, the sarcomere organization of the RRC-1::HA strain is also normal. Next we used HA antibodies to perform immunostaining to localize RRC-1::HA in muscle. As demonstrated in Figure 7b, these antibodies localize RRC-1-HA to the MCB, co-localizing with antibodies to PAT-6 (α-parvin). We also observed weak localization of RRC-1-HA to the same focal plane as the base of the M-lines and dense bodies where PAT-6 is localized, in a generally striated pattern, but in a more diffuse, less organized manner. This weak staining is not background staining, as the same dilution of anti-HA antibodies and the same gain with our confocal microscope detects no fluorescence in wild type animals (Figure 7b, left column). The pattern of RRC-1-HA is somewhat similar to the pattern of UNC-52 (perlecan) immunostaining (Qadota et al., 2017). These results show that RRC-1 is localized to MCBs, and to the bases of M-lines and dense bodies which is consistent with RRC-1 playing a role in the formation or stability of these structures and the structure of the sarcomere.

**Figure 7.**
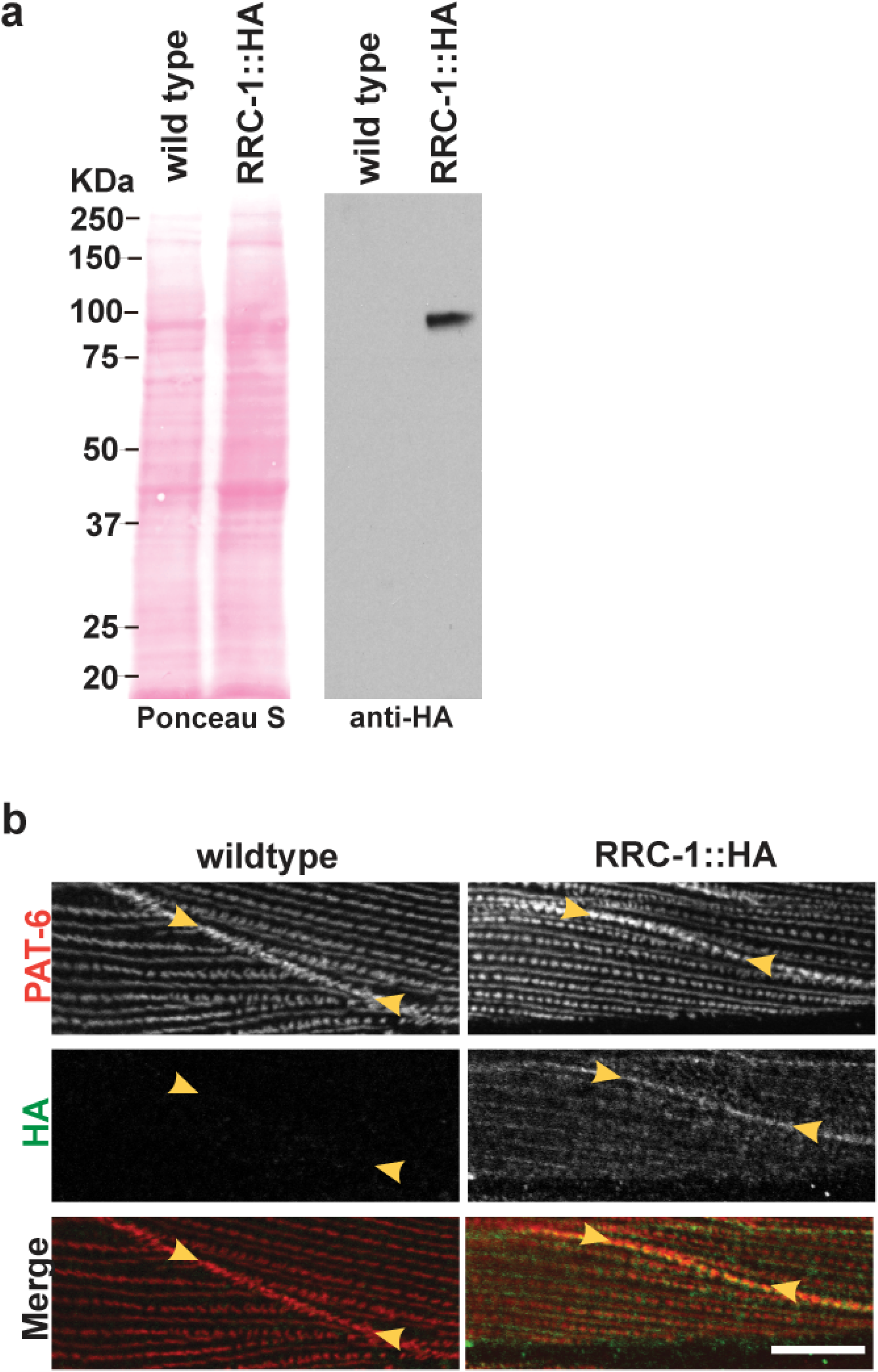
HA-tagged RRC-1 localizes to muscle cell boundaries. **a**.) Confirmation that the CRISPR/Cas9-generated strain, *rrc-1(syb4499)* expresses RRC-1::HA. Lysates were prepared from wildtype and *rrc-1(syb4499)*, and portions separated by SDS-PAGE, blotted, and reacted with antibodies to HA. Anti-HA detects a protein of expected size, approximately 90 kDa, from *rrc-1(syb4499)* and not from wild type. **b.)** Confocal microscopy imaging of body wall muscle co-stained with anti-PAT-6 (α-parvin) and anti-HA antibodies in wildtype and the CRISPR generated strain that expresses RRC-1::HA. Note that RRC-1 localizes to the muscle cell boundary, co-localizing with PAT-6. There is weaker localization of RRC-1::HA to the same focal plane at the bases of M-lines and dense bodies, in a striated pattern, but in a diffuse, less-organized manner. Scale bar, 10 μm.

### Genetic interaction between *rrc-1*, *pix-1* and *git-1*

Given the similar localization of RRC-1 and PIX-1 to MCBs, dense bodies and M-lines, and that mutants in either gene affect MCB organization, we sought to determine if there are genetic interactions between these two genes. By genetic recombination, we created a strain that expresses the HA tagged RRC-1 in a *pix-1* mutant background, which we designate, RRC-1::HA *pix-1(gk299374)*. We compared the immunolocalization of PAT-6, and RRC-1::HA in a wild type vs. the *pix-1* mutant background. As shown in Figure 8a, PAT-6 is normally localized to the MCB in the strain expressing RRC-1::HA, but missing in the strain RRC-1::HA *pix- 1(gk299374)*, as expected for this *pix-1* mutant (Moody et al., 2020). When we immunostained with anti-HA, we found that RRC-1::HA is mostly missing from the MCB in RRC-1::HA *pix- 1(gk299374)*. However, the total protein level of RRC-1::HA is not affected by *pix-1* deficiency, as shown by a quantitative western blot (Figure 8b and c). These data suggest PIX-1 is required for RRC-1 localization to the MCB.

**Figure 8.**
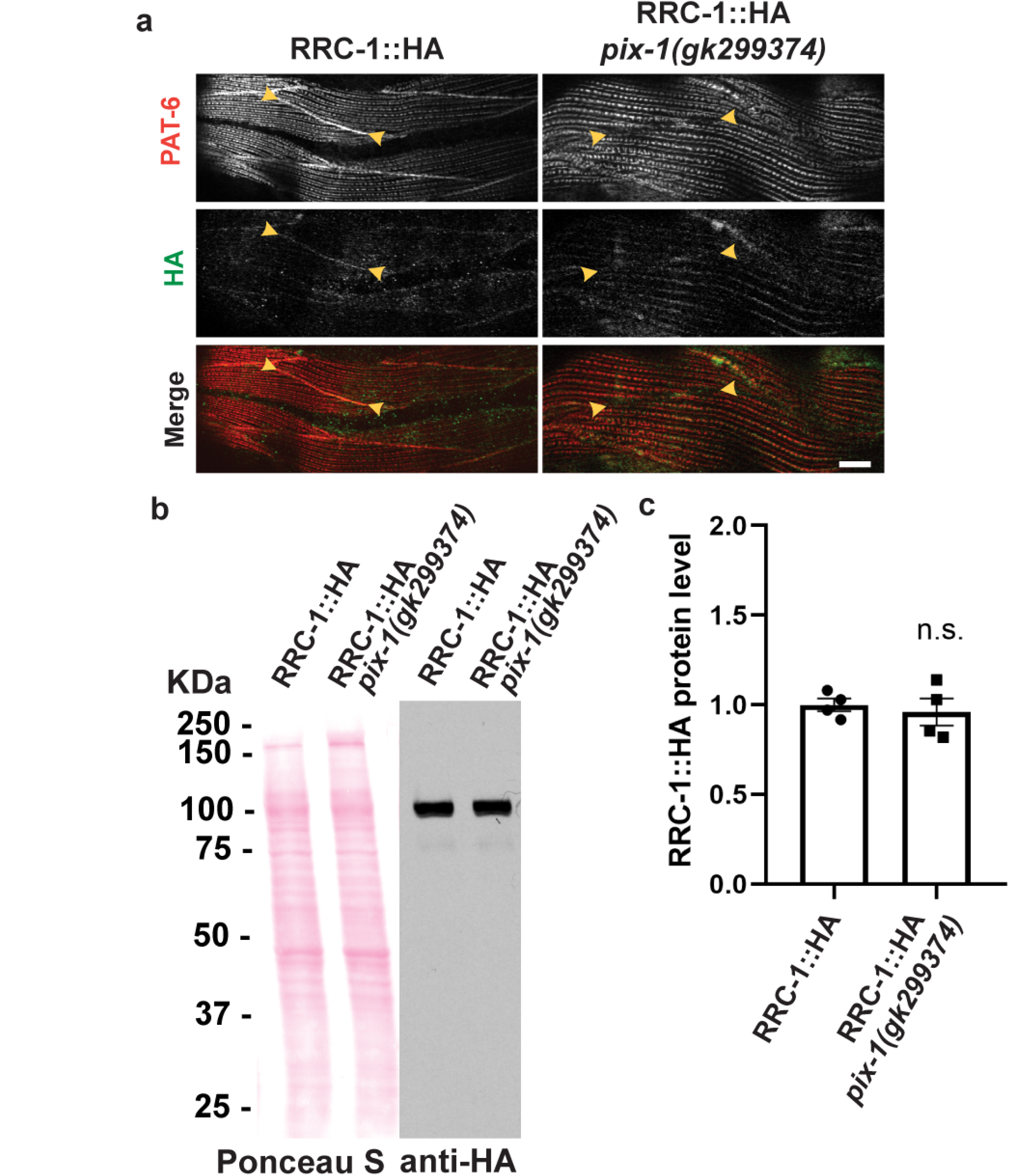
PIX-1 is required for the proper localization of RRC-1 but not the stability of RRC-1. **a.)** Confocal microscopy imaging of body wall muscle co-stained with anti-PAT-6 (α-parvin) and anti-HA antibodies in the CRISPR generated strain that expresses RRC-1::HA, and in a strain that expresses RRC-1::HA in a *pix-1* null background. Note that when PIX-1 is deficient there is less localization of RRC-1::HA to the muscle cell boundary. Scale bar, 10 μm. **b.)** Western blot showing that the level of RRC-1::HA is the same in wild type vs. the *pix-1* mutant. **c.)** Quantification of HA-tagged RRC-1 protein levels in wild type vs. *pix-1* mutant shows no significant (n.s.) difference using a Welch’s t-test for statistical analysis, N=4.

We next asked whether *rrc-1* deficiency could affect the localization of PIX-1. We compared the localization of PIX-1 in wild type vs. *rrc-1(ok1747)* and *rrc-1(tm1023)*. As shown in Figure 9a, there is much less accumulation of PIX-1 at MCBs in the *rrc-1* mutants as compared to wild type. However, the total level of PIX-1 is not affected (Figure 9b and c). Therefore, localization of PIX-1 to the MCB depends on RRC-1. Finally, we asked whether GIT-1 is required for the localization or stability of RRC-1. Previously, we reported that GIT-1, a scaffold for PIX-1, is required for the assembly or stability of IACs at the MCB, and showed that GIT-1 is required for the stability of PIX-1, by using a *git-1* deletion allele, *git-1(ok1848)*(Moody et al., 2020). We now show that *git-1* is also required for the localization of PIX-1 to the MCBs, as shown in Figure 10a. To examine the localization of RRC-1-HA in a *git-1* mutant required a recombinant, but because the two genes are so close together on the X chromosome (<0.5 cM apart), we employed *git-1* RNAi, instead. As shown in Figure 10b, knock down of *git-1* results in a reduced level of PAT-6 at MCBs, as expected. *git-1* RNAi also results in RRC-1-HA still being localized at MCBs but perhaps at a slightly reduced level. Nevertheless, *git-1* RNAi does result in a significant reduction in the level of RRC-1-HA by western blot (Figure 10c and d). As indicated in Figure 10d, the level of RRC-1-HA is reduced by four-fold compared to empty vector. Overall, our results indicate that PIX-1 is required for the proper localization of RRC-1 but not its stability, that RRC-1 is required for the localization of PIX-1 but not its stability, and that GIT-1 is not required for the localization of RRC-1 but is required for its stability.

**Figure 9.**
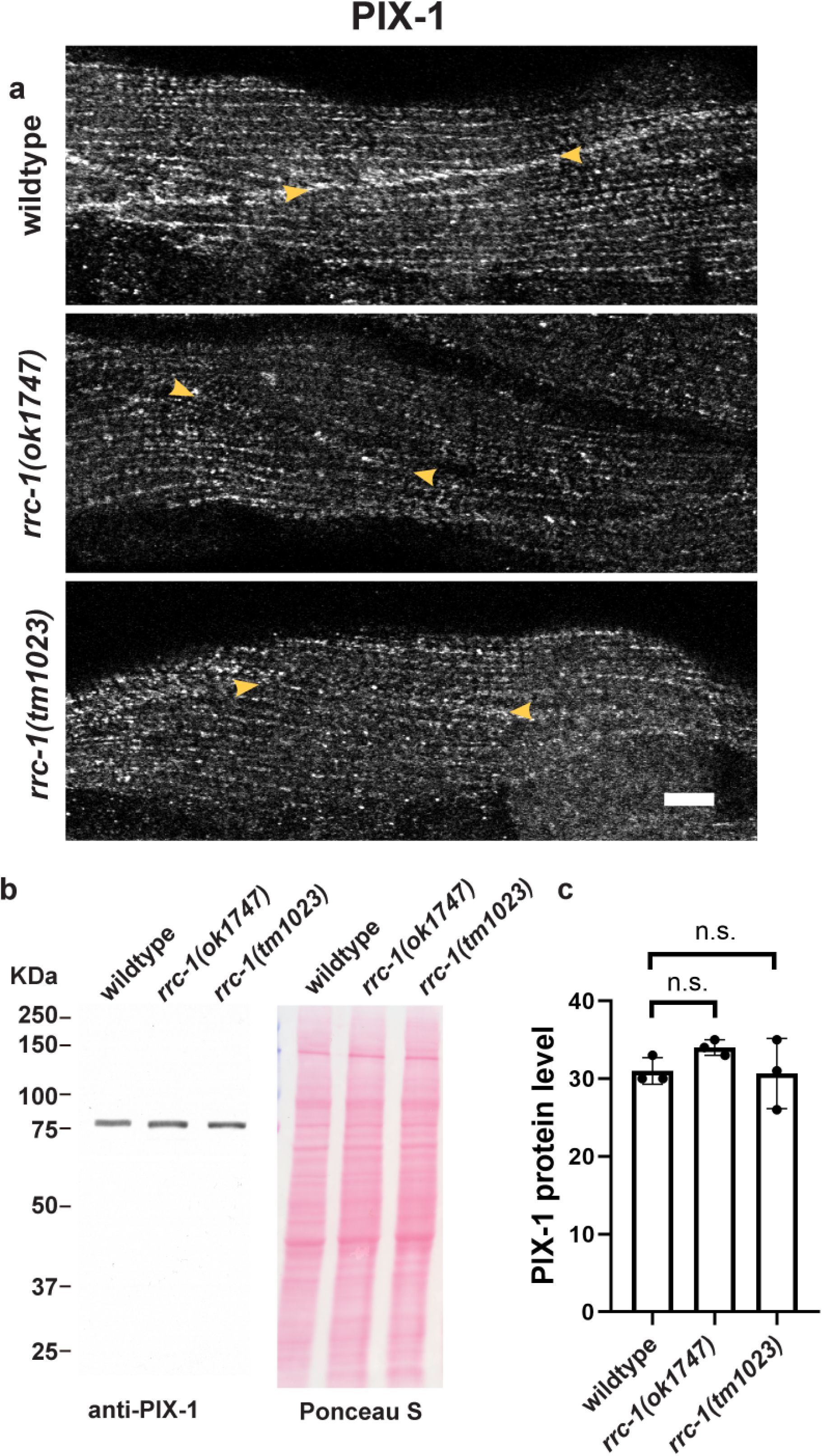
RRC-1 is required for the proper localization of PIX-1 but not the stability of PIX-1. **a.)** Confocal microscopy imaging of body wall muscle stained with antibodies to PIX-1 in wild type and two deletion alleles of *rrc-1*. Note that when RRC-1 is deficient the localization of PIX- 1 to the muscle cell boundaries is nearly absent (indicated by yellow arrowheads). Scale bar, 10 μm. **b.)** Western blot showing that the level of PIX-1 is the same in wild type vs. the *rrc-1* mutants. **c.)** Quantification of PIX-1 protein levels in wild type vs. either *rrc-1* mutant showing no significant (n.s.) differences using a Welch’s t-test for statistical analysis, N=3.

**Figure 10.**
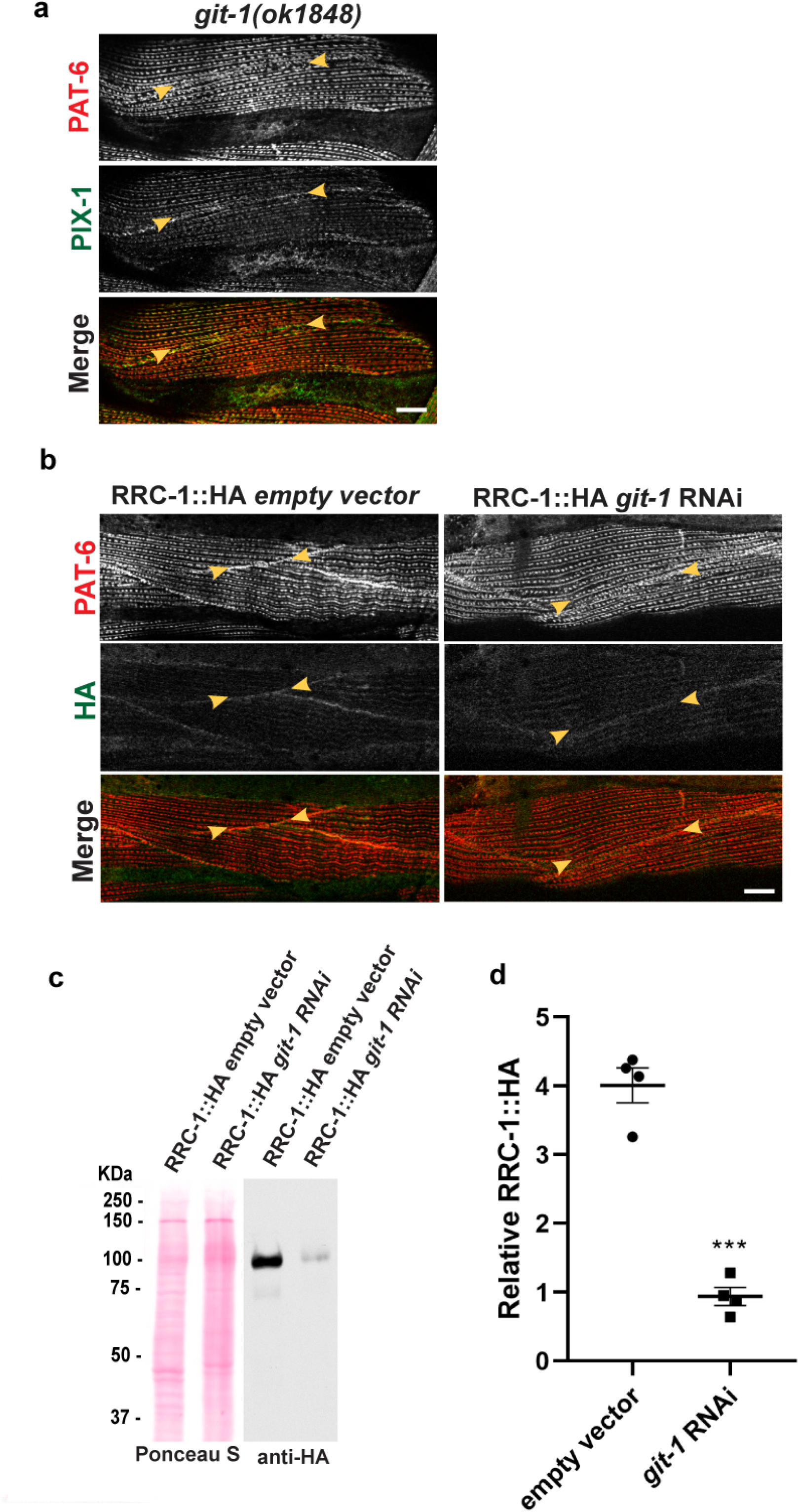
GIT-1 is not required for the localization of RRC-1, but GIT-1 is required for the stability of RRC-1. **a.)** Confocal microscopy imaging of body wall muscle co-stained with anti-PAT-6 (α-parvin) and anti-PIX-1 antibodies in wildtype and in the *git-1(ok1848)* deletion mutant. Note that both PAT-6 and PIX-1 show poor localization to the muscle cell boundaries. Scale bar, 10 μm. **b.)** Confocal microscopy imaging of body wall muscle co-stained with anti-PAT-6 (α-parvin) and anti-HA to detect RRC-1::HA in a strain expressing RRC-1::HA with and without feeding bacteria expressing dsRNA for *git-1*. Note that RNAi knockdown of *git-1* does not prevent the localization of RRC-1::HA to the muscle cell boundaries. Scale bar, 10 μm. **c.)** Western blot showing the level of RRC-1::HA in wild type vs. *git-1(RNAi)*. Note that *git-1(RNAi)* results in a reduced level of RRC-1::HA. **d.)** Quantification of HA-tagged RRC-1 protein levels in *git-1 (RNAi)* vs. empty vector control shows that *git-1 (RNAi)* reduces the level of RRC-1::HA to about 23% of the level in wild type. A Welch’s t-test for statistical analysis, N=4, was used; Error bars indicate SEM, ***: p≤ 0.001.

Given that GIT-1 is required for the stability of RRC-1, we explored whether RRC-1 might be in a protein complex with GIT-1. To address this question, we used CRISPR/Cas9 to add a mNeonGreen tag to the C-terminus of GIT-1 in the strain PHX4499, which expresses RRC-1 with a C-terminal HA tag. The resulting strain, called PHX5908, expresses both RRC-1-HA and GIT-1-mNeonGreen. A total worm lysate of PHX5908 was made, and using nanobodies to mNeonGreen coupled to magnetic beads, we were able to immunoprecipitate (IP) GIT-1- mNeonGreen, running at approximately 120 kDa (Figure 11). This IP was then examined by western blot for coIP of RRC-1-HA, PIX-1 and actin (as negative control). As shown in Figure 11, we were able to detect coIP of PIX-1, consistent with the reported high affinity binding between PIX and GIT proteins in mammals (Schlenker and Rittinger, 2009). In addition, what was coIP’d appears to be specific, as actin was detected in the lysate (L) but not the IP. Unfortunately, although RRC-1-HA could be detected in the lysate (L), even after a long exposure, we could not detect coIP of RRC-1-HA.

**Figure 11.**
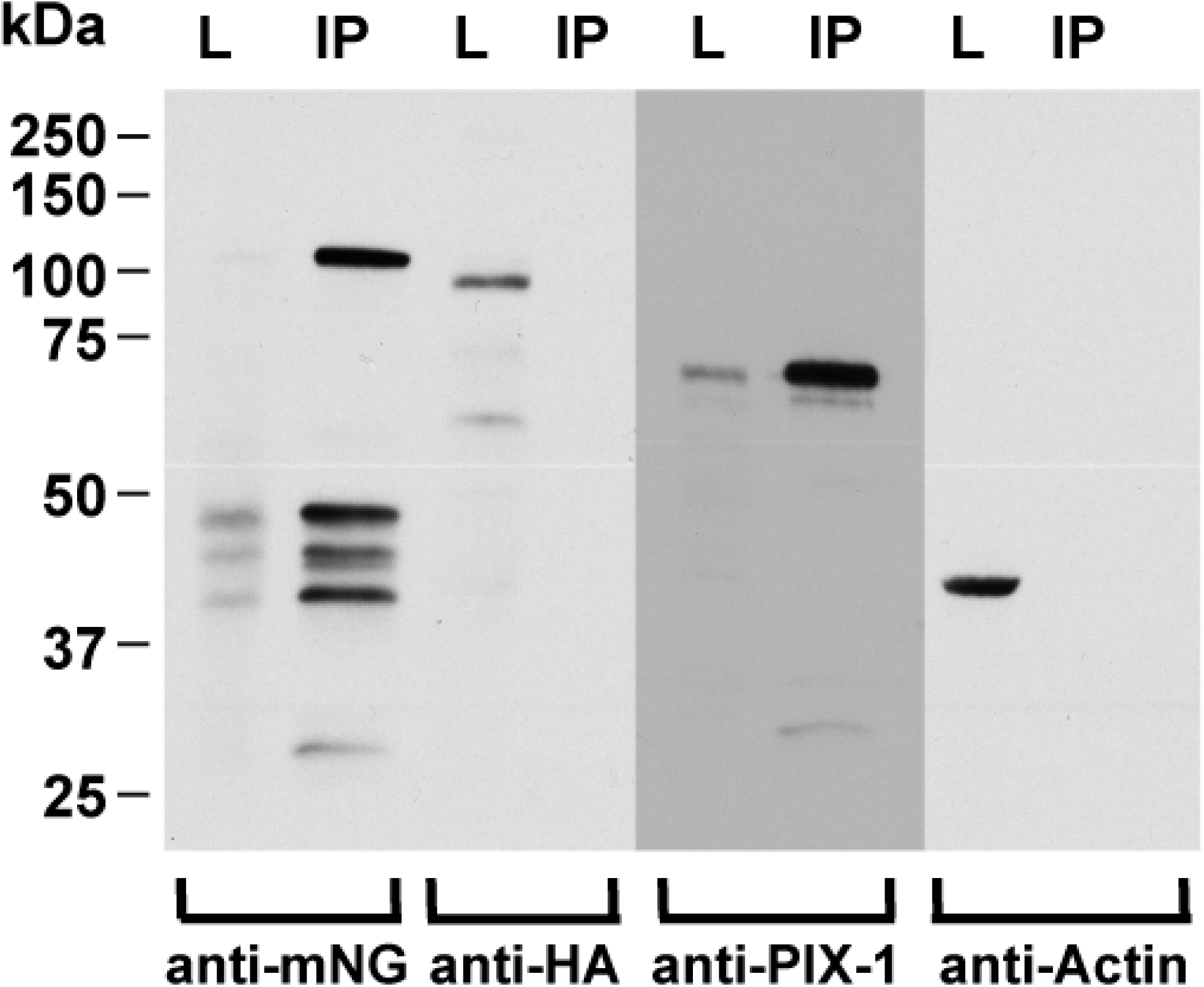
Co-IP analysis of GIT-1. GIT-1-mNeonGreen was immunoprecipitated from a CRISPR-generated worm strain that expresses both GIT-1-mNeonGreen and RRC-1-HA. Portions of the total worm lysate (L) and immunoprecipitated (IP) were separated by SDS-PAGE, transferred to membrane, and reacted with the indicated antibodies. As shown from left to right, the IP procedure worked well, and clearly pulled down GIT-1-mNeonGreen from the lysate, a major band of expected size (∼100 kDa), as well as some bands 40-50 kDa that are likely degradation products. Although detected in the lysate (L), no detectable RRC-1-HA was detected in the IP. Nevertheless, the IP does contain PIX-1, which is clearly concentrated in the IP vs. the L. (In fact, the anti-PIX-1 reaction was so strong, that a lower exposure to film is shown (the reason for the change in background). In addition, the IP was fairly specific: although actin can be detected in L, no actin was detectable in the IP.

## Discussion

The studies reported here identify RRC-1 as a Rho family GAP required for the assembly or stability of IACs in *C. elegans* muscle, and probably acting at least partially in the PIX-1 pathway. Loss of function of *rrc-1* results in disorganization of the IACs at all of its locations in nematode muscle—MCBs, M-lines and dense bodies (Figures 2 and 4). A likely consequence of this IAC disorganization is the disorganization of the sarcomeres and reduced muscle function (locomotion) observed in *rrc-1* mutants. (Figures 5 and 6). Our conclusion that RRC-1 functions as a GAP for the PIX pathway are based upon the following: (1) Loss of function for mutations in the GEF, PIX-1, and the GAP, RRC-1, each reduce the accumulation of IAC components at the adhesion plaques of the MCB. That loss of function of a GEF, PIX-1, a positive regulator of Rac, and loss of function of a GAP, RRC-1, a negative regulator of Rac, have the same defect at MCBs, suggests that the maintenance of integrin adhesion complexes at MCBs is a dynamic process. Consistent with these results, if we increase or eliminate the protein kinase activity of a known effector of the PIX-1 pathway, PAK-1, we also observe defects in the MCB (Figure 1). (2) RRC-1 and PIX-1 each localize to the MCB (Figure 7). (3) RRC-1 localization to the MCB requires PIX-1 (Figure 8). (4) PIX-1 localization to the MCB requires RRC-1 (Figure 9). (5) GIT-1, a scaffold for assembly of PIX-1 and PAK-1, when knocked down by RNAi results in a reduced level of RRC-1 (Figure 10). (6) Both PIX-1 and RRC-1 affect the activity of Rac: we have reported that a *pix-1* null mutant has reduced levels of activated GTP-bound CED-10 (Rac) in nematode muscle (Moody et al., 2020). By expressing nematode RRC-1 in mammalian tissue culture cells, Delawary et al. (2007) have reported that RRC-1 has GAP activity towards mammalian Rac and Cdc42 but not RhoA. Figure 12 summarizes what we now know about the *pix-1* pathway in *C. elegans* body wall muscle. Of course, we still do not know the substrates for PAK-1 and PAK-2 protein kinases in muscle and how their phosphorylation results in the assembly or maintenance of IACs at the muscle cell boundary.

**Figure 12.**
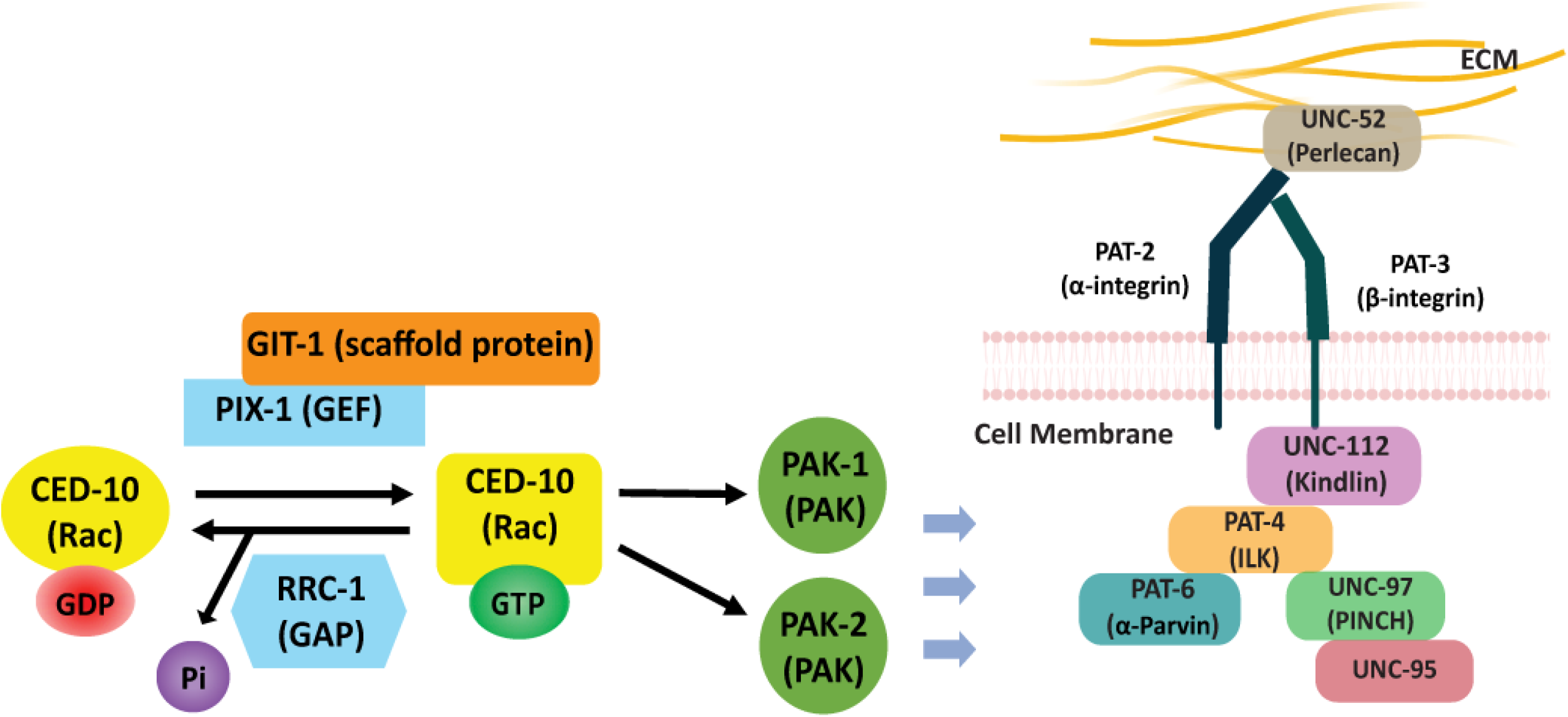
RRC-1 is Rho GAP in the PIX-1 pathway. The drawing depicts what we have learned about the PIX-1 pathway in *C. elegans* muscle. In Moody et al. (2020) we demonstrated that for the proper assembly or stability of integrin adhesion sites at the muscle cell boundary, the Rac GEF, PIX-1, its scaffold, GIT-1, the Rac, CED-10, and the PAK effectors, PAK-1 and PAK-2 are required. The current results are most consistent with the Rac GAP, RRC-1 being the GAP for the PIX pathway in striated muscle, based on the similarity of *pix-1* and *rrc-1* phenotypes, and genetic interactions of *rrc-1* with *pix-1* and *git-1*. The blue arrows indicate that, by some still unknown mechanisms, this Rac pathway is required for the assembly or stability of integrin adhesion complexes at muscle cell boundaries. Insight into these mechanisms likely will emerge by identifying the substrates for PAK-1 and PAK-2 in nematode muscle, which are currently unknown. Core components of this complex are depicted to the right, with the names of the mammalian proteins shown in parentheses.

We wondered if RRC-1 might exist in a complex with GIT-1. This was suspected because *pix-1* and *rrc-1* mutants similarly affect the MCB; PIX-1 and RRC-1 have similar intracellular localizations; GIT is a known scaffold for PIX and PAK in mammals; and because we showed in this study (Figure 10) that RNAi knock down of *git-1* results in reduced levels of RRC-1. We could IP GIT-1 and demonstrate coIP with PIX-1, consistent with the known high-affinity binding between GIT and PIX proteins in mammals, but we were unable to detect coIP with RRC-1 (Figure 11). There are several possible reasons for this outcome including that the conditions we used to solubilize GIT-1 destroyed interaction with RRC-1, that we did not solubilize enough RRC-1, or that the tags included—mNeonGreen and HA, each included at the C-termini of GIT-1 and RRC-1, respectively—interfered with the interaction. The last possibility seems unlikely as RRC-1-HA is properly localized to the MCB in the strain that expresses both RRC-1-HA and GIT-1-mNeonGreen (data not shown). Unfortunately, using anti-mNeonGreen antibodies, we could not detect localization of GIT-1-mNeonGreen in muscle. The PIX pathway has been shown to be functionally important in multiple organisms and tissues, ranging from mammalian nervous (Ramakers et al., 2012) and immune (Missy et al, 2008) systems, to nematode germline (Lucanic and Cheng, 2008), migration of neuroblasts (Dyer et al., 2010), tension-dependent morphogenesis of epidermal cells (Zhang et al., 2011), early embryonic elongation (Martin et al., 2014), and muscle (Moody et al. 2020). Given that we have identified two possible human orthologs for RRC-1, ARHGAP32 and ARHGAP33 (Figure 3), and each are known to be expressed in skeletal and heart muscle, we propose that one or both of these proteins are GAPs for the PIX pathway in human muscle. As noted above, in *C. elegans*, *pix-1* has also been shown to function in neurons. During the migration of nematode QR neuroblast descendants, noncanonical signaling through PIX-1 occurs (Dyer et al. 2010), but when migration stops, canonical Wnt signaling is engaged by activating EVA-1 in the Slt-Robo pathway, and the RhoGAP called RGA-9 (Rella et al., 2021). However, note that PIX-1 functions in activation of neuroblast migration, but RGA-9 functions to inhibit neuroblast migration. Therefore, RGA-9 might be a RhoGAP for the PIX-1 pathway in neuroblast migration, but this is difficult to conclude from the existing data. Also note that although *rga-9* is expressed in muscle, we did not detect a MCB defect in a deletion allele of the gene (Supplementary Figure 1).

In addition to *rrc-1*, our screening of mutants in 18 Rho GAP proteins expressed in muscle revealed two other genes that when mutated result in MCB defects, *hum-7* and *rga-4* (Supplementary Figure 1). *hum-7* mutants only affect the MCB (like *pix-1* mutants), whereas the *rga-4* mutant affects the MCB, M-line and dense body (like *rrc-1*). *hum-7* encodes a 213 kDa protein with Rho GAP domain near its C-terminus and a myosin class IX motor domain in its N-terminal half. Inspection of two independently generated mutants have the same MCB-specific defect as *pix-1*. However, we have only examined a single mutant allele of *rga-4*, the intragenic deletion *rga-4(ok1935)*, and it had not been outcrossed to wild type. *rga-4* encodes a 1126 aa protein with the only recognizable domain being the Rho GAP domain. RGA-4 has been reported to act redundantly with another Rho GAP protein, RGA-3, in the germ line and early embryo, and to inactivate RHO-1 (RhoA) of *C. elegans* (Schmutz et al., 2007). Interestingly, we have previously reported that RNAi knockdown of RHO-1 (RhoA) results in disorganization of the A-bands in the body wall muscle of adult nematodes (Qadota et al., 2008). We leave investigation of *hum-7* and *rga-4* in muscle for future studies.

## Materials and Methods

### *C. elegans* strains

All nematode strains were grown on NGM pates using standard methods and maintained at 20° (Brenner, 1974). Most strains were obtained from the Caenorhabditis Genetics Center. The wild type strain was N2 (Bristol). Strains containing mutations in 18 genes encoding proteins with Rho GAP domains and also expressed in muscle, are listed in Supplementary Table 1 and in Supplementary Figure 1. The following strains were generated during this study:

GB340: *pak-1(syb647)*, which contains a L99F mutation in PAK-1, was generated by CRISPR/Cas9 by SunyBiotech (described below) as PHX647, and then outcrossed 4X to wild type.

GB341: *pak-1(syb632)*, which contains a K324A mutation in PAK-1, was generated by CRISPR/Cas9 by SunyBiotech (described below) as PHX632, and then outcrossed 1X to wild type.

GB342: *rrc-1(syb4499)*, which expresses RRC-1 with an HA tag fused to its C-terminus (RRC- 1::HA), was generated by CRISPR/Cas9 by SunyBiotech (described below) as PHX4499.

GB343: *rrc-1(ok1747)* was outcrossed 5X to wild type.

GB344: *rrc-1(tm1023)* was outcrossed 5X to wild type.

GB345: *rrc-1(gk290525)* was outcrossed 5X to wild type

GB346: *rrc-1(gk859353)* was outcrossed 5X to wild type

GB348: *rrc-1(syb4499) pix-1(gk299374)* was generated by recombination starting with GB342 and GB291 (Moody et al., 2020).

DM8008: raIs8 [unc-112::GFP + rol-6(su1006)]

### CRISPR/Cas9 generation of nematode strains expressing kinase dead and kinase constitutively-active PAK-1, HA-tagged RRC-1, and mNeonGreen-tagged GIT-1

The CRISPR/Cas9 procedures were carried out by SunyBiotech (http://www.sunybiotech.com). Details about the sgRNAs and repair templates used are given in Supplementary Figure 5. The resulting strains are:

PHX647, *pak-1(syb647)* which has an L99F mutation predicted to make the PAK-1 protein kinase constitutively active

PHX632, *pak-1(syb632)* which has a K324A mutation predicted to make the PAK-1 protein kinase catalytically dead

PHX4499, *rrc-1(syb4499)* which expresses RRC-1 with a C-terminal HA tag. PHX5908, *rrc-1(syb4499) git-1(syb5908)* which expresses both RRC-1::HA and GIT- 1::mNeonGreen.

### Immunostaining and confocal microscopy of body wall muscle

Adult worms were fixed and immunostained using the method described by Nonet et al. (1997), and additional details described by Wilson et al. (2012). Antibodies were used at 1:200 dilution except as noted: anti-PAT-6 (rat polyclonal)(Warner et al., 2013), anti-UNC-52 (mouse monoclonal MH2)(Mullen et al., 1999), anti-UNC-95 (rabbit polyclonal Benian-13)(Qadota et al., 2007), anti-UNC-112 (1:100 dilution)(Hikita et al., 2005), anti-MHC A (mouse monoclonal 5–6)(Miller et al., 1983), anti-UNC-89 (rabbit polyclonal EU30)(Benian et al., 1996), anti-ATN-1 (mouse monoclonal MH35)(Francis and Waterston, 1991), and anti-HA (rabbit monoclonal C29F4 from Cell Signaling Technology). Secondary antibodies, used at 1:200 dilution, included anti-rabbit Alexa 488, anti-rat Alexa 594, and anti-mouse Alexa 594, all purchased from Invitrogen. Fixation and phalloidin-rhodamine staining was conducted as described (Waterston et al., 1984). Images were captured at room temperature with a Zeiss confocal system (LSM510) equipped with an Axiovert 100M microscope and an Apochromat x63/1.4 numerical aperture oil immersion objective, in 1× and 2.5× zoom mode. For all the confocal images the color balances were adjusted by using Adobe Photoshop (Adobe, San Jose, CA).

### Live imaging of UNC-112-GFP

We live imaged UNC-112-GFP using the integrated transgenic array strain DM8008 (raIs8 [unc- 112::GFP + rol-6(su1006)]) which expresses full-length UNC-112 with a GFP insertion. To determine the localization of UNC-112-GFP in *pix-1(gk299374)* and *rrc-1(ok1747)*, we crossed DM8008 into these *pix-1* and *rrc-1* mutants. The transgene was followed by using the Rol-6 marker, and the presence of the *rrc-1* or *pix-1* mutations was followed by PCR and/or DNA sequencing. Live imaging was performed as described in Moody et al. (2020). Essentially, animals were washed off plates using M9 buffer and immobilized with 10 μM levamisole in M9 for 10 min. Approximately 50-100 animals in 3 μl were added to 7 μl of ice-cold 25% Pluronic F127 in M9 (Hwang et al., 2014) lying on a cold glass slide, to which was added a cover slip and sealed with nail polish. After incubation at room temperature for 5-10 min to solidify, images were taken using a confocal microscope as described above.

### Swimming and crawling assays

For swimming assays day two asynchronous adults were harvested from one 6 cm NGM OP50 seeded plate with M9 buffer. Animals were subsequently washed free from bacteria using M9 buffer and then pelleted at ratio of 1:1 (worm: buffer). 2 ml of M9 buffer followed by five microliters of worm suspension were added to the center area of one unseeded 6 cm NGM plate. Each strain was allowed to adapt for five minutes before recording swimming movement. The recordings were done using a dissecting stereoscopic microscope fitted with a CMOS camera (Thorlabs). For all strains a total of fifteen, 10-sec. videos were recorded from various sections of the plate with each video tracking an average of 8 individual animals. Video data was analyzed by Image J WrmTracker software plug-in to obtain body bends per second (BBPS) for individual animals. Worms that moved out of frame and outliers were removed during data analysis and an average of 20 animals were analyzed for each strain. The resulting BBPS values for each mutant strain was compared to wildtype and further tested for statistically significant differences using Welch’s T-test.

For crawling assays day two adults were harvested as described above, except for the use of 0.2g/L gelatin in M9 buffer. Five microliters of worm suspension was added to the center of a 6 cm unseeded NGM plate and then the excess liquid was removed using a twisted KimWipe. After a five-minute adaptation time, worm crawling movement was recorded as mentioned above. BBPS values for individual animals were extracted from each video. The resulting values for each strain was compared to wild type for statistical analysis using Welch’s T-test for significance.

### Protein sequence analysis

Nematode RRC-1a, b, and c were obtained from Wormbase. By using a BLAST homology search, we confirmed that ARHGAP31, ARHGAP32 and ARHGAP33 are the human proteins most homologous to RRC-1. The domain organization for RRC-1 and its homologs were analyzed by the online PFAM database. PubMed pBLAST database was used to align human ARHGAP31-33 amino acid sequences with nematode RRC-1 to determine the percent identities for each domain and total protein.

### Knockdown of GIT-1 via RNAi feeding

RNAi by feeding was performed as described previously (Timmons et al., 2001; Miller et al. 2009). GIT-1 cDNA was generated via PCR amplification of the 5’-most 1,077 nucleotides of the GIT-1 cDNA sequence using the RB2 cDNA library as a template with the following primers--GIT-1 FWD: 5’GCGGGATCCATGTACACAGCAGAGGCGCTT 3’which includes a BamHI restriction enzyme (RE) site and GIT-1 REV: 5’CGCCTCGAGTGCTGGATTGTCTCCAGTGAT 3’, which includes an XhoI RE site. The ∼1kb amplicon was digested and ligated into the BamHI and XhoI sites of the pPD129.36 vector, and used to transform competent XL1 Blue E. Coli cells on LB + ampicillin plates overnight at 37 °C. Individual colonies from the GIT-1 cDNA pPD129.36 clones were grown overnight in liquid culture, plasmids prepared, and confirmed by restriction digestion. A resulting GIT-1 pPD129.36 clone and empty vector pPD129.36 plasmids were used to transform competent HT115 (DE3) RNAi feeding bacteria, and a resulting colony from each was grown as a overnight liquid culture. The resulting bacteria were used to seed 6cm and 10 cm NGM plates. To conduct RNAi feeding experiments, 15-20 L4 stage worms were added to 25 NGM *git-1* RNAi and 25 empty vector in HT115 (DE3) bacteria 6cm plates and left overnight. Then following day, ten worms were transferred from the 6cm plates to the 10cm plates under the same conditions and allowed to lay eggs for approximately 8 hrs before being picked. The eggs on the plate were allowed to hatch for ∼48 hrs before being harvested for conducting fixation for immunostaining or making lysates for SDS PAGE, followed by Western blotting analysis. (Miller et. al 2009)

### Western blot analysis

The method of Hannak et al. (2002) was used to prepare total protein lysates from wild-type, *rrc- 1(ok1747)* 5X O.C., *rrc-1(tm1023)* 5X O.C., RRC-1::HA, RRC-1::HA *pix-1(gk299374)*, RRC-1::HA; RNAi empty vector, and RRC-1::HA; *git-1 (RNAi)* mixed-stage animals. Equal amounts of total protein were separated on 10% polyacrylamide-SDS- Laemmli gels, transferred to nitrocellulose membranes, reacted with affinity purified, E. coli-OP50-absorbed anti-PIX-1a (Moody et al., 2020), anti-HA (rabbit monoclonal cat. no. C29F4 from Cell Signaling Technology) at 1:1,000 dilution, and anti-histone H3 (rabbit polyclonal ab1791, Abcam, Inc.; 1:40,000 dilution), then reacted with goat anti-rabbit immunoglobulin G conjugated to HRP (GE Healthcare) at 1:10,000 dilution, and visualized by ECL. Protein bands were quantitated by comparing to total Ponceau S staining or to histone H3.

### Immunoprecipitation of GIT-1::mNeonGreen

A large quantity of worms (∼3.5 ml) from strain PHX5908 (*rrc-1(syb4499) git-1(syb5908*)) which expresses both RRC-1-HA and GIT-1-mNeonGreen, were grown on 20, 15 cm-high peptone NGM plates seeded with E. coli strain NA22, and a “worm powder” was generated by grinding the worms extensively in a mortar and pestle in liquid nitrogen. A total protein lysate was prepared by adding worm powder to 1 ml of “RIPA Buffer” (∼20% vol/vol) consisting of 50 mM Tris pH 7.5, 150 mM NaCl, 1% Nonidet P-40, 0.5% sodium deoxycholate, 0.1% sodium dodecyl sulfate, 1 mM EDTA, and cOmplete Mini protease inhibitor (Roche), vortexing for 1 minute, incubating on a rotating wheel at 4° for 30 minutes, vortexing for 1 minute, spinning at top speed in a microfuge at 4° for 10 minutes, and saving the supernatant. 100 μl of supernatant was concentrated and desalted using an Amicon Ultra-0.5 centrifugal filter device according to the manufacturer’s instructions (Millipore, Inc.), re-constituted in an equal volume of 50 mM Tris 7.5, 1 mM EDTA, and after adding an equal volume of 2X Laemmli sample buffer, mixing and heating to 95°, this material was designated as lysate (L). We found that this desalting step was necessary to avoid distortion of an SDS-PAGE upon electrophoresis. To the remainder of the supernatant was added 30 μl of mNeonGreen-Trap Magnetic Agarose, which are nanobodies to mNeonGreen covalently attached to magnetic beads (cat. no. ntma-20; Chromotek, Inc.), incubating on a rotating wheel at 4° for 1 hour, removing the beads from the solution using a rack containing neodymium magnets, and then washing the beads with 1 ml of Wash Buffer (50 mM Tris pH 7.5, 150 mM NaCl, 1 mM EDTA, cOmplete Mini protease inhibitors) 3X. To the washed beads was added 36 μl of 2X Laemmli sample buffer, vortexing for 5 seconds, heating at 95° for 5 minutes, vortexing for 5 seconds, and then separating out the beads on the magnetic stand; the resulting liquid was designated as “immunoprecipitate” (IP). Multiple lanes containing either 12 ml of L and IP, were separated on a 10% SDS-PAGE, transferred to nitrocellulose membrane, and as shown in Figure 10, portions of the blot were reacted against anti-mNeonGreen (mouse monoclonal antibody 32F6 from Chromotek, Inc., at 1:200 dilution), anti-HA (rabbit monoclonal antibody, cat. no. 3724 from Cell Signaling, Inc., at 1:1,000 dilution), anti-PIX-1 (rabbit polyclonal antibody (Moody et al., 2020) at 1:500 dilution), and anti-actin (mouse monoclonal C4 cat. no. MA5-11869 from Invitrogen, Inc., at 1:5,000 dilution), reacted with the appropriate HRP-conjugated secondary antibodies and visualized with ECL.

## Supporting information

Supplemental Figure 1 part 1

Supplemental Figure 1 part 2

## Acknowledgments

We thank SunyBiotech Corporation for generation of the CRISPR/Cas9 strains PHX632, PHX647, and PHX4499, Sophia Figueroa and Jordan Walter for generating a full-length cDNA for *rrc-1*, Robert Barstead (Oklahoma Medical Research Foundation) for the cDNA library RB2, and Andrew Fire (Stanford University) for the RNAi vector pPD129.36. We thank Richard Kahn and Dorothy Lerit for helpful comments on the manuscript. Most of the nematode strains used in this work were provided by the Caenorhabditis Genetics Center, which is funded by the NIH Office of Research Infrastructure Programs (P40 OD010440). This study was supported in-part by an NSF-Graduate Research Fellowship (DGE 1444932) to J.C.M., and from a NIH grant (R01HL160693) to G.M.B.

## Author contributions

J.C.M. helped design and interpret the experiments, carried out nearly all of the experiments, and helped write the paper. H.Q. helped design the experiments, helped with his knowledge of the topic, and helped write the discussion. G.M.B. helped design and interpret experiments, carried out a few of the experiments, and wrote the paper together with J.C.M and H.Q.

## Conflict of Interest

The authors declare no competing or conflicts of interest.

## Supplementary Material

**Supplementary Table 1.** Rho GAP Screening Results in tabular form.

**Supplementary Figure 1. Rho GAP Screening Results.** Confocal microscopy imaging of body wall muscle from mutants in 18 genes that encode proteins harboring Rho GAP domains that are also expressed in muscle, immunostained with antibodies to PAT-6 (α-parvin). Yellow arrowheads indicate the disrupted muscle cell boundaries in *rrc-1*, *hum-7* and *rga-4* mutants. Scale bar, 10 μm.

**Supplementary Figure 2. UNC-112::GFP is present at the muscle cell boundary of wild type but is missing at the muscle cell boundary in *pix-1* and *rrc-1* mutants.** Representative confocal images of body wall muscle from strains expressing UNC-112::GFP from an integrated transgenic array in wild type, *pix-1* and *rrc-1* mutants. Yellow arrowheads indicate the positions of muscle cell boundaries. Scale bar, 10 μm.

**Supplementary Figure 3. Addition of an HA tag to the C-terminus of RRC-1 has no effect on whole worm locomotion.** Top, swimming assays; bottom, crawling assays. For each assay, there is no apparent difference in the distribution or means of wild type vs. the strain expressing RRC-1::HA. For each strain, N=20 for the swimming assays, and N=16 for the crawling assays. Welch’s t-test was used to test for significance. Error bars indicate SEM. n.s.: no significant difference.

**Supplementary Figure 4. Addition of an HA tag to the C-terminus of RRC-1 has no effect on sarcomere organization.** Confocal images of PHX4499 which expresses RRC-1::HA reacted with phalloidin-rhodamine (thin filaments), and antibodies to sarcomere proteins MHC A (thick filaments), UNC-89 (M-lines) and ATN-1 (dense bodies). Each image is a representative image obtained from at least two immunostaining experiments. Scale bar, 10 μm.

**Supplementary Figure 5. sgRNAs and repair template information for generation of CRISPR/Cas9 strains.**

## Notes

### Competing Interest Statement

The authors have declared no competing interest.

### Summary of Updates

We have confirmed a muscle cell boundary defect in rrc-1 defects by live imaging of UNC-112-GFP (Supplemental Figure 2). More careful study revealed that in addition to defects at the muscle cell boundary, in he more severe alleles, the M-lines and dense bodies are disorganized, and that the sarcomere is also disorganized (Figure 5). rrc-1 mutations result in mislocalization of PIX-1 (Figure 9).

