## Supplementary figures and images for "The Rho GAP RRC-1 is required for the assembly or stability of integrin adhesion complexes and is a member of the PIX pathway in muscle"

### Supplemental Figure 1 part 1

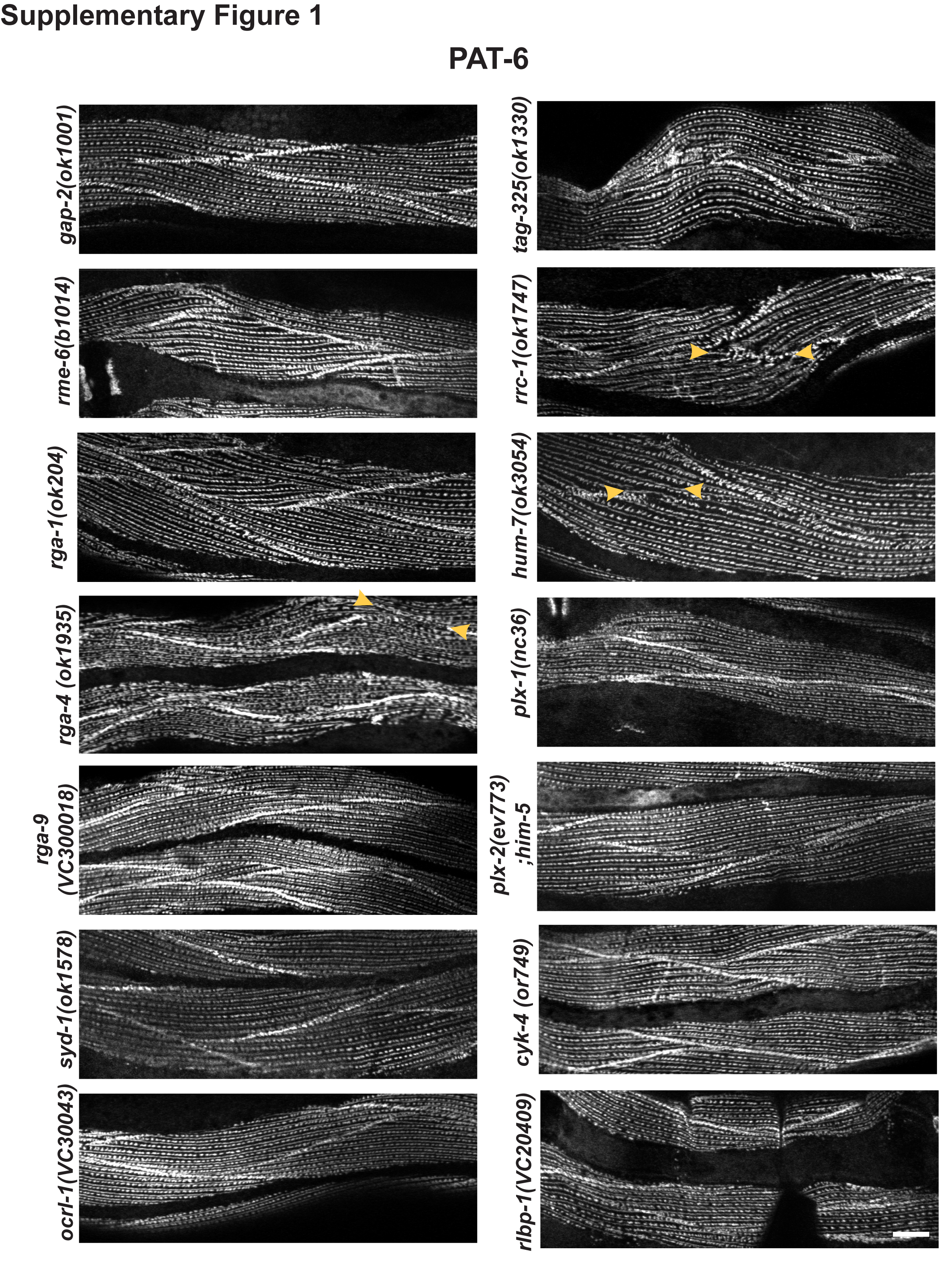

### Supplemental Figure 1 part 2

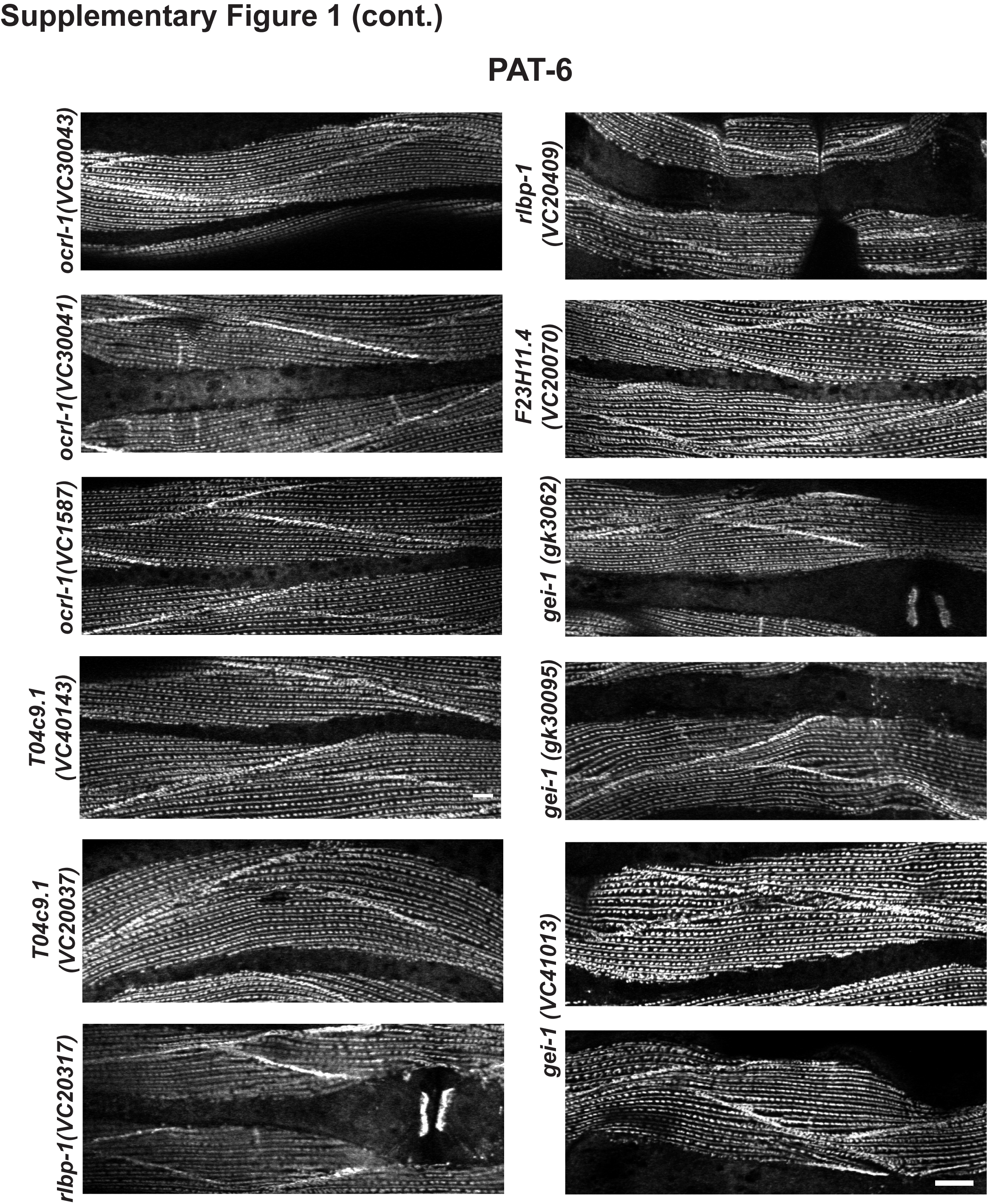

### Supplemental Figure 2

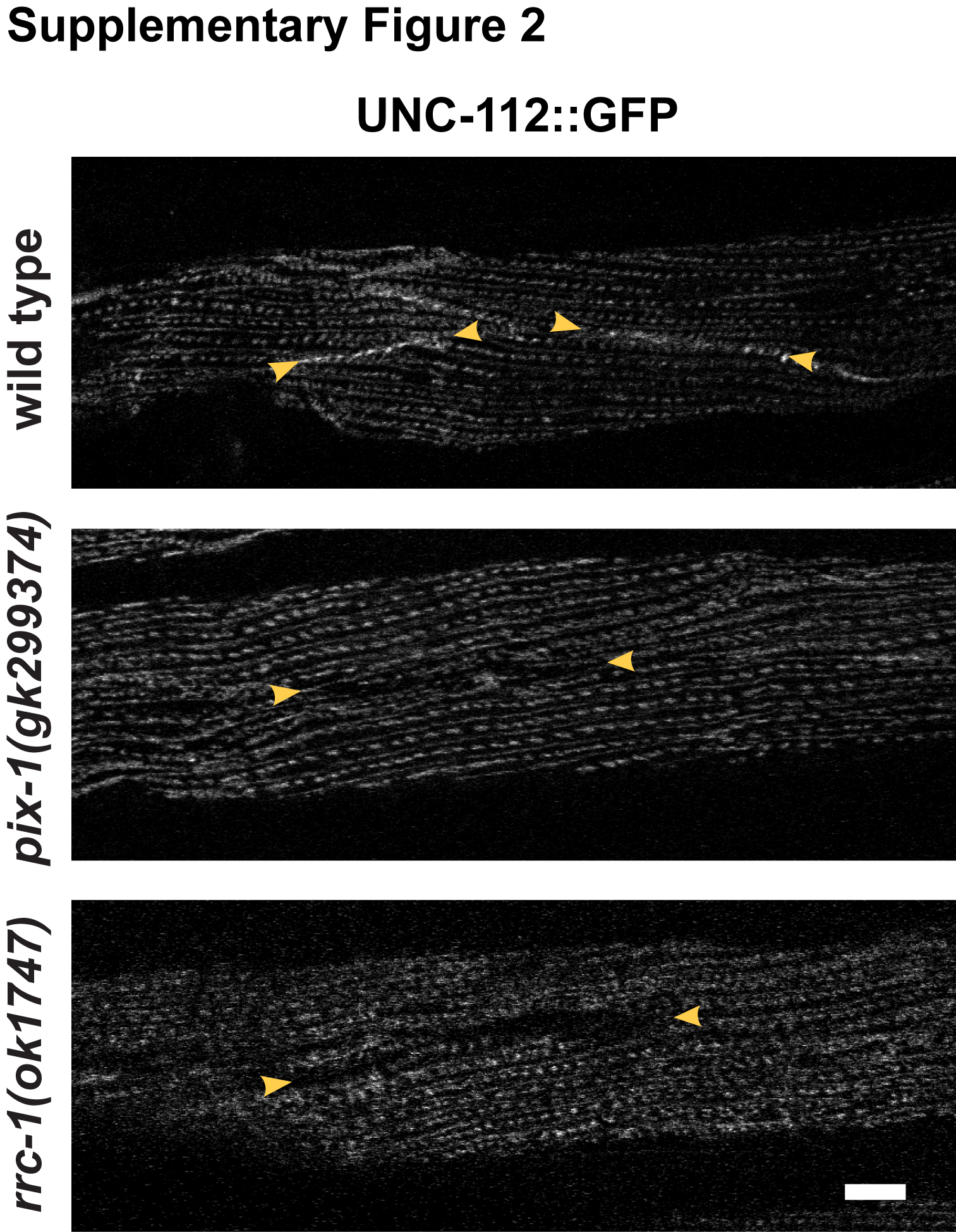

### Supplemental Figure 3

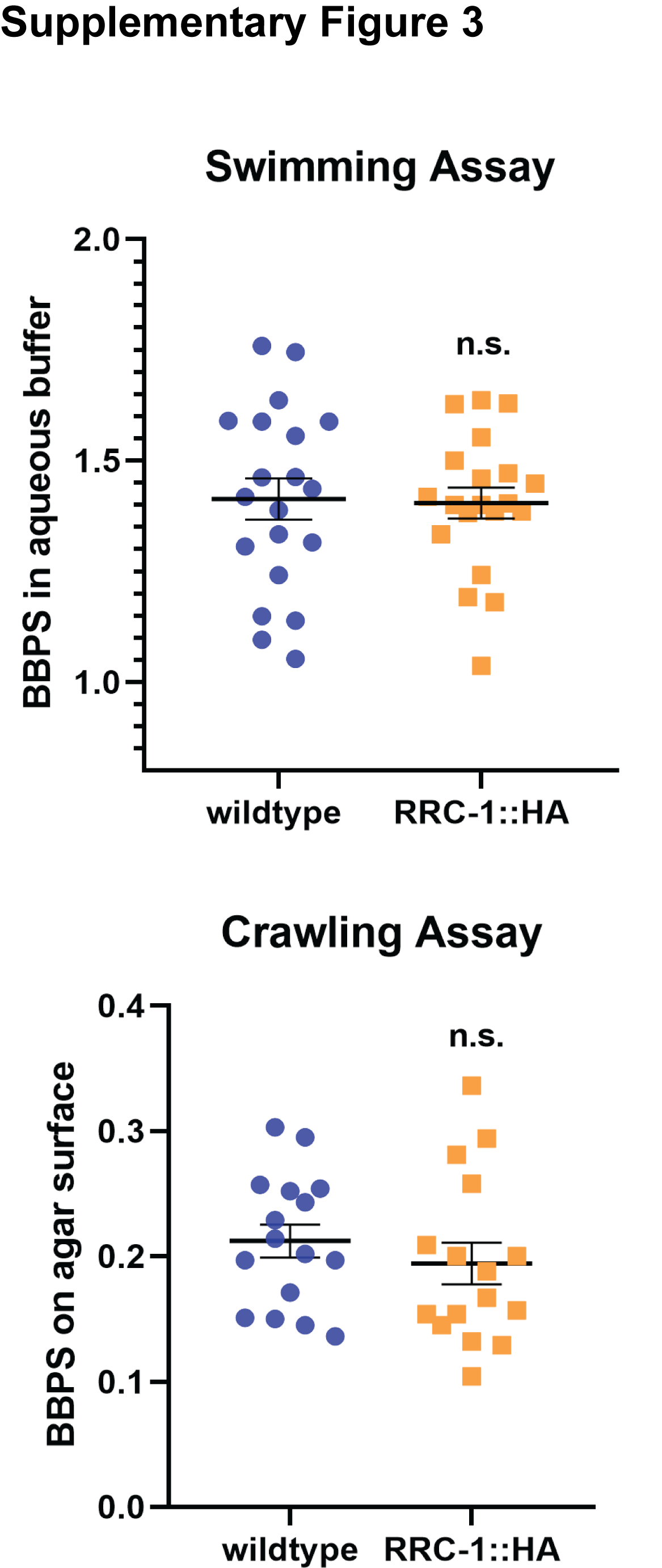

### Supplemental Figure 4

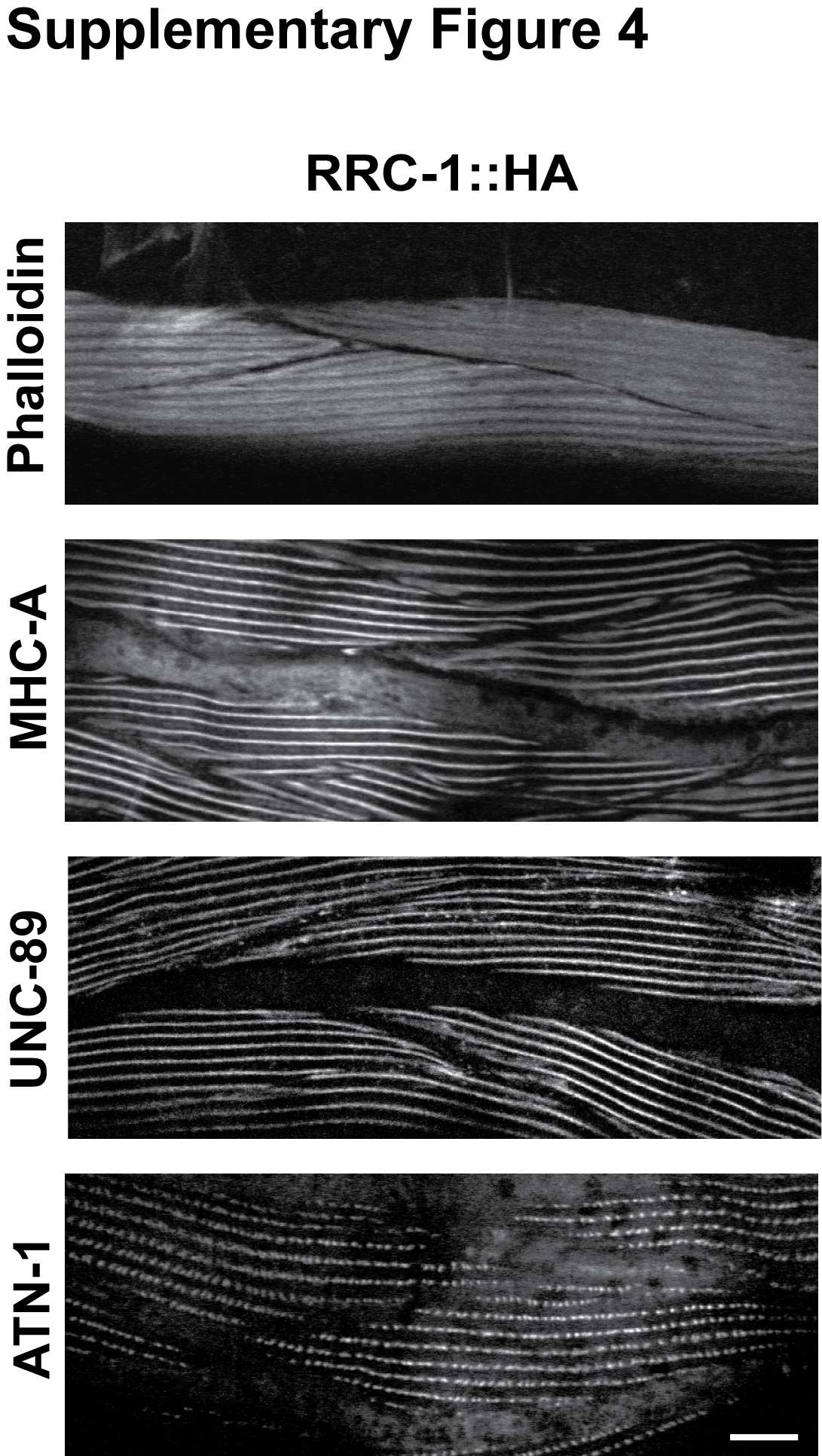
