## Supplemental Figure 5 for "The Rho GAP RRC-1 is required for the assembly or stability of integrin adhesion complexes and is a member of the PIX pathway in muscle"

**Supplementary Figure 5**

Information on generation of CRISPR/Cas9 strains

Synonymous mutations are labeled in blue font

* PAM sites are marked by squares

**PHX632:**

Sg1: CCGAAGCGGAGGTGGCTATCAAG

Sg2: TATTGAAATTAGTACCGAAGCGG

**>Repair template**

tacttgtgtattacatgcccttagttttagagtatttttttaagttaaaaacaataaaaacgtgtaccacattttttatttctaggtttaggaaataatctttctaagcctggagaggaaaaatgtatatcacgtttaaattatatggagaaatgaatgaaaatttaattcaatttcaaaatgagggcatgtaatacacaagtaccgaatttttcccaagttctctataacatataagtatattttcagTACACAAAACCGAAAGAGGAGGAAGAAAAAATTCCAGACCTTTCAAAAGGACAATTTGGTGTACAGGCCAGAGGTCAAAAAGCTAAGAAAAAGATGACTGACGCTGAAGTGCTGACTAAGCTCCGTACCATTGTGTCTATCGGAAATCCAGATCGAAAATATAGAAAAGTTGATAAAATCGGCTCAGGTGCATCTGGTTCTGTGTACACCGCTATTGAAATTAGT**ACG**GAAGCGGAGGTGGCT**ATTGCT**CAGATGAACCTGAAGGATCAACCAAAGAAGGAATTGATCATTAATGAGATTTTGGTGATGCGTGAGAATAAGCATGCAAATATTGTAAATTATTTGGATTCGTATTTGGTGTGCGATGAATTATGGGTAGTGATGGAGTATCTTGCCGGTGGATCATTGACTGATGTTGTCACGGAGTGCCAGATGGAGGATGGAATTATTGCAGCTGTTTGCAGAGAAGTTCTTCAAGCGCTTGAATTCCTCCACAGCCGCCACGTCATTCACAGAGATATTAAATCTGACAATATTCTTTTGGGAATGGATGGTTCGGTGAAATTGAgtaagattataatttttaatgcttttgcaacctacttctgtgaatttcagCCGACTTTGGATTCTGTGCTCAGCTCTCGCCGGAGCAAAGAAAACGCACGACAATGGTCGGAACTCCATACTGGgtgagtgatggaaattgaaaaagcgaataggaaattaatgtttgacaattgcagATGGCGCCGGAAGT

**PHX647:**

sg1: CCTGAAGCATGGGCACGTCTTCT

sg2: CCTCAGGCAGTGTTGGACGCGCT

**>Repair template**

ttctgacaccggtcgcaaccgctaatcgtaccagattaacgtgaatgctaatagtaagctaaaaactaaaaactactacttttactttttccctcagtcaaaaaatcattataaacaaatctcaatttataataatattttgcaaccgttaattcagtgcattatcttctaaaaacgtgttattcttgaaaccatggttagtgttaataaagcttgcgtttttgagacgttttaagaaaacaaatccatacgagaaaaatttaagtgaagccaattatcacgaacatgtattcaaccaaagatagtttcttctaaagacctatgaaattgtcccaatttttttagtatgaagaggtctgttttgtcgaatagacttgagaaactactgtagaatcttatttataaacttttctcagaaacttttgctccttatcaacaatataaattttcagGGAATGCCT**GAG**GCATGGGCACGT**TTGTTT**ACAGACTCACAGATCTCAAAACAAGAGCAGCAACAGAATCCT**CAA**GCAGTGTTGGACGCGCTCAAATACTACACACAAGGCGAAAGCAGCGGCCAGAAGTGGTTGCAGTACGATATGAgtaagtaacccgtagtcagtcttgctaatggagctcattcttctacagTGTTTATAGATGACGCACCTTCTCGGACGCCATCATACGGACTGAAACCGCAACCATATAGCACATCATCCCTGCCGTATCATGGCAATAAAATTCAGGATCCAAGAAAGATGAATCCAATGACAACCAGTACAAGTAGTGCGGGGTATAACAGCAAGCAAGGAGTTCCTCCGACGACGTTTAGTGTAAATGAGAATAGATCGAGTATGCCACCGgtaagtgggaagaataatcttgaagttcatattgattgcttcagAGTTATGCACCGCCACCGGTCCCCCATGGTGAAACTCCTGCTGATATTGTTCCTCCCGCTATCCCTGATAGGCCGGCAAGGACGTTGAGTATTgtgagtagaatttttgg

**PHX4499:**

sg1: CCGATAACTAACCTTTCTCCAGC

sg2: CCTTTCTCCAGCTGCACAAATGC

sg3: CCAGTCATTTGTTCTCTGCATAA

**>Repair template**

aattctgaaatacaggttgtaaaaaaaagataatggcaaccatcttatttttcagAACACGTGGCAACATTCCATGAAAGATCGAGTCCTGTAGAAGAATGGTCAAGTGATTCTAGAGAGAGTCTTCATCTTGAAATGTCCCGTTATGATAACGTATCTCCAAGTGGTACTATTACAAGAAgtgagtttaaaaaaaaaaatttatgacattaacaaatatgtagaaagcagtctttcttttattgtatttcaaatgggtaacagcaaaattactttaatacaaagtttcaaagtcaataatccacagcaaattcaaacatttcttgtggttttttgatctgtacaaaataaatcttagccaaatttggtcaatttccagccacaaacactataatttcacaataacacaataattcttggcgtttcagATCAACGAGAACCGATAACTAAC**tTg**TCTCCAGCTGCACAAATGCTCTTTTTCGAATCTTCTCGA**GCg**AGTCATTTGTTC**AGC**GCATACCCATACGATGTTCCAGATTACGCTGGTGGATCTTACCCATACGATGTTCCAGATTACGCTGGTGGATCTTACCCATACGATGTTCCAGATTACGCTTAAttgaatttccacctctttcatgatatttttgttcattgtattgttgactactttttcaactttatttctttatcgtcttaaatttttaaagaaccagtgttccattttttcatttcctcatatttttcttctacttctcaattgtcttcaagcaattcctgatcatttttattttgtttcgtgtttttgattataaattatttatatacaaacacacaattttcaaacaattgtgatgcgtgataacatcgctgaattgtatttttgaaattataaaagaaacgcggaatacaagacctaattactagaatttactgaatttaaataaacactttttagtgatttatttcgttttgaaaattttccgcgatcctgatttgaagaattataacaatttcaatcaattctaatttttattttcaatattttgttcaaagttcaaaacgatattgtggagttgtaaaaaaaattaaaatatatattcaaatatgcattacat

PHX5908: (note: CRISPR/Cas9 editing was carried out on strain PHX4499)

BG38-out-s:CCTTGCAGGCTAGTTCATAG

BG38-out-a:TTCCTCCATTCTGACGGTTG

BG38-seq-s:ATTTTGCTCCCGTACGTGAG

BG38-seq-a:CACACGTTCCGAACCTTTCA

>Repair template

ccttgcagGCTAGTTCATAGCTTACAAGCCCAAAACATCAATCTCAACTCGGAGCTGATCACATTCAGAGATGACTTGTACAATTTGAAGAGAACAAATATGACAAGAAGGATGCCATCTCCATTGGCTATTGTAGAGACTCCAGAAAGAGTTATGCCACCAGTGGGAATTCAACAAGGTTTTTCAAGGAGgtaaaacaaaaaacaaattcagatatttttacaaaatttttgaagttgacaagttctttggaatttcatttccatctcctaatatgtaagactacaagtcagggatgcgcgacattcgaacgttttaagcaataagcaatgttgacaactgctagttttaataattgccgagcagcaattgccgcgcacccctactacaaatattttttaatattttcatgacttacatacatttggttgattaaaatgttctactattgtttaatattcagAGATAGTGAAGATAGAATCGATGGTGGCGGAGCTCGGTGGAGGTCAAATTCCAGTGACCGTAGAAACTCAAACGACCACAAACTGGACAAAAAAGTAGAAAGACGTCAGGACTCAATGATGGAAAGCTCGTCTTCCCTATCACGCTCCTCACAAAACCCATCTTGCCTCAATAGAGAAGgtaggattaacccgcaacgtctttccctcagttcatccatgttttcagAGGTGAAAACTCGCGTTATTCAACAAAGTGAAAAAATCACGCGCCACATTAAAACCCTGCTGCAACACGGACATAATGGAAACCTAGACATGAACGCACGCGGCGGTGCACACGACgtaagtttggacagctgaaacgaagtacgaaaacttgtcatttttgcagATTGCTTGCGCCATCAATAGTCTTATCACTATTTTGCTCCCGTACGTGAGACACGAGAAAATCGATACTTTGACGGACGCCGTGGTGgtaagtagaagaacaaatgttgaacttcgtgaattaagtagtagattaaatcaatgttcaatacatactgcattttttctattagtcttgcacctgaaggggccaatcaaaaaatcattgtatcgcagcctctgatagacttacacaagctttttgccagaggtttgctgtcttgtagtctctattaatcttgtgcccctacggttcaattaaaaattagtcttgcatgcaagcctaataagggagacacggtatttataatcgtgaatataaatatttattgcttaaatttcaaatctaaaatttagCTTTTGAATGCAAAATGCAATAGCCCAGTG**CTT**ATGCCAATGGATATTGTCGACGCAGCACAAACCATTGCTGAAAAACTTCGC**CTCATC**ATCATGGAATTTTGTATGGTCTCCAAGGGAGAGGAGGACAACATGGCCTCCCTCCCAGCCACCCACGAGCTCCACATCTTCGGATCCATCAACGGAGTCGACTTCGACATGGTCGGACAAGGAACCGGAAACCCAAACGACGGATACGAGGAGCTCAACCTCAAGTCCACCAAGgtaagtttaaacatatatatactaactaaccctgattatttaaattttcagGGAGACCTCCAATTCTCCCCATGGATCCTCGTCCCACACATCGGATACGGATTCCACCAATACCTCCCATACCCAGACGGAATGTCCCCATTCCAAGCCGCCATGGTCGACGGATCCGGATACCAAGTCCACCGTACCATGCAATTCGAGGACGGAGCCTCCCTCACCGTCAACTACCGTTACACCTACGAGGGATCCCACATCAAGgtaagtttaaacagttcggtactaactaaccatacatatttaaattttcagGGAGAGGCCCAAGTCAAGGGAACCGGATTCCCAGCCGACGGACCAGTCATGACCAACTCCCTCACCGCCGCCGACTGGTGCCGTTCCAAGAAGACCTACCCAAACGACAAGgtaagtttaaacatgattttactaactaactaatctgatttaaattttcagACCATCATCTCCACCTTCAAGTGGTCCTACACCACCGGAAACGGAAAGCGTTACCGTTCCACCGCCCGTACCACCTACACCTTCGCCAAGCCAATGGCCGCCAACTACCTCAAGAACCAACCAATGTACGTCTTCCGTAAGACCGAGCTCAAGCACTCCAAGACCGAGCTCAACTTCAAGGAGTGGCAAAAGGCCTTCACCGACGTCATGGGAATGGACGAGCTCTACAAGTGAtttctcttcatcctagaatgctttttctttattatatatattatatatcttaattataattacctgttttgttaattattcagttggtttttttcttgctcaatttttatacattgtttaagcaataagtgtaatttaaaagtgtcttatttattagcatttaaatgtcgattaccatttgattttttttctcattgttggcatatttcatgacttttcgctctcattgcttcaccaaaaattttcgtctcgtcttgaacaaaaactgaaacataataaacttaatgaaaaacatgccttatataaattttagttttagagaagtcacctagtgcaaagtgaagtgaaaggttcggaacgtgtgaaaaattatgtacaaggtttaagaaggttttaatttcatcagtttttttttggcatgataggggaacaagttcaaagtagaaaaatctaagtattatccaccttttaattgtggtcaaacaaatacatttgaatcaaaaattactcacaatgaatctaaagacaaaatttatatgtgtagacaagtaaaaactatgtacttaccatttggttaaatttaaggctctttgaaaaacattaataaaccgctaaggtctctacgttcgaaaacactgactaaatcttatcccaatatatgcgaatagggaactactattctaatcttaaataatttcaaattaatatttcacctagataggttacactactggtttcgtaaatttgaagcacaataaaagaagaacaaagttttgcacggtggccttttcgtcgttttatctttaatattacttcacagattacctctcagcaactggcaaatacattcttctgtgactgaaaaaaagccgctgccatctccacagaatcatcttttcttttcaatttcatttttcgagcgactgataaggaaggtgcgatagtgtcggtttaggggtgttaagggaattggcaagtgaatgagcactgagcaaagcaattaaaagtttagcgacaacgacctggagaggtgtgcgatcccacagtttagcggagagacagatcaaactatttgatggaagcgagggtaacgccaaaaaagtcataaaggtgaatatgcaaagaaggaaaagcggctgataagaaacagagcgctgttgaaactcatttgcgtttccccttttttgtttgctgcgatttgtcttgtccctcacaccctcatcatcaactggtacacgaccgcattttctgccatttcatttttctccagtatttgtaaaaagaatgaccattgagagagctactcgtcaaaatgttcgaagaggacgaccaccaggtacacgaagagcagttgtttttgatgacacaccacaaccgtcagaatggaggaa
