## Supplemental Table 1 for "The Rho GAP RRC-1 is required for the assembly or stability of integrin adhesion complexes and is a member of the PIX pathway in muscle"

### Supplementary Table 1: RhoGAP Screening Results.

| Genes with RhoGAP domains | Muscle Expression | MCB phenotype |
|---|---|---|
| CHIN-1 | No | No |
| CYK-4 | Yes | No |
| F23H11.4, isoform a | Yes | No |
| GAP-1 | No | No |
| GAP-2, isoform a | Yes | No |
| GAP-3, isoform a | No | No |
| GEI-1, isoform a | Yes | No |
| HUM-7, isoform a | Yes | Yes |
| OCRL-1, isoform a | Yes | No |
| PAC-1, isoform a | No | No |
| PES-7 | No | No |
| PLX-1, isoform a | Yes | No |
| PLX-2 | Yes | No |
| RGA-1, isoform a | Yes | No |
| RGA-2 | yes | No |
| RGA-3 | No | No |
| RGA-4, isoform a | Yes | Yes* |
| RGA-5, isoform b | No | No |
| RGA-6, isoform a | No | No |
| RGA-8, isoform a | No | No |
| RGA-9, isoform a | Yes | No |
| RLBP-1, isoform a | Yes | No |
| RME-6, isoform a | Yes | No |
| RRC-1, isoform a | Yes | Yes |
| SPV-1, isoform a | No | No |
| SRGP-1, isoform a | No | No |
| SYD-1, isoform a | Yes | No |
| T04C9.1, isoform a | Yes | No |
| TAG-325, isoform a | Yes | No |
| Y92H12BL.4 | No | No |

*: M-lines and dense bodies are also disorganized
